# Topographically distinct adaptive landscapes for teeth, skeletons, and size explain the adaptive radiation of Carnivora (Mammalia)

**DOI:** 10.1101/2022.04.01.486739

**Authors:** Graham J. Slater

## Abstract

Models of adaptive radiation were originally developed to explain the early, rapid appearance of distinct modes of life within diversifying clades. Phylogenetic tests of this hypothesis have yielded limited support for temporally declining rates of phenotypic evolution across diverse clades, but the concept of an adaptive landscape that links form to fitness, while also crucial to these models, has received more limited attention. Using methods that assess the temporal accumulation of morphological variation and estimate the topography of the underlying adaptive landscape, I found evidence of an early partitioning of mandibulo-dental morphological variation in Carnivora (Mammalia) that occurs on an adaptive landscape with multiple peaks, consistent with classic ideas about adaptive radiation. Although strong support for this mode of adaptive radiation is present in traits related to diet, its signal is not present in body mass data or for traits related to locomotor behavior and substrate use. These findings suggest that adaptive radiations may occur along some axes of ecomorphological variation without leaving a signal in others and that their dynamics are more complex than simple univariate tests might suggest.

## Introduction

Modern theories of morphological evolution often make reference to the idea of an adaptive landscape. The topography of the adaptive landscape relates gene frequencies (Wright, 1932) or, less strictly, morphological variation (e.g., Martin and Wainwright, 2013) to fitness, and provides a mechanism for understanding evolutionary change within populations at the generational scale (Lande, 1976; Arnold et al., 2001; Uyeda et al., 2011). G. G. Simpson was the first to make the crucial connection between models of the adaptive landscape taken from population genetics and macroevolutionary patterns observable in the fossil record, leading him to propose that rapid phenotypic evolution in small, peripherally isolated populations (Mayr, 1942) provided a mechanism for the geologically sudden appearance of distinct adaptive types without long-lived intermediate forms via a process he called quantum evolution (Simpson 1944, see also Grant 1963). In later linking quantum evolution to the phenomenon of adaptive radiation (Osborn, 1899, 1902), Simpson (1953) hypothesized that this mode of diversification would likely be concentrated early in clade history when peaks on the adaptive landscape, which he called adaptive zones, were vacant. As adaptive zones fill and ecological opportunity declines, the rate at which novel phenotypes are produced should subsequently slow (Osborn, 1902; Simpson, 1953; Valentine, 1980; Walker and Valentine, 1984), yielding a predictable signal in functional trait data.

One challenge that macroevolutionary biologists face is to infer the topography of the adaptive landscape, and historical dynamics on it, from comparative morphological data. The development of statistical approaches for evaluating tempo and mode in phenotypic evolution on phylogenetic trees has resulted in a rich and diverse toolkit for this task. Some authors, noting that Simpson’s model implies that rates of phenotypic evolution should decline over clade history as adaptive zones fill, have used phenomenological (Harmon et al., 2003) or explicit models based on time-varying rates (Blomberg et al., 2003; Freckleton and Harvey, 2006; Harmon et al., 2010) to evaluate support for an “early burst” pattern in comparative data. Relaxing the requirement that shifts to new adaptive zones are concentrated early in clade history has further resulted in models that permit shifts in evolutionary rates as a function of co-occuring species diversity (Mahler et al., 2010) or along branches of a phylogeny (Venditti et al., 2011; Eastman et al., 2011; Rabosky, 2014) and jumps in trait means independent of evolutionary rate variation (Landis et al., 2013; Landis and Schraiber, 2017; Duchen et al., 2017; Pagel et al., 2022). Other authors have noted that adaptive evolution towards optimal phenotypes implies very different long-term dynamics and covariances between species than a purely diffusive process (Hansen and Martins, 1996a), leading to a variety of methods to infer the presence and location of shifts toward new trait optima or, more precisely, the topography of the underlying adaptive landscape, based on a mean-reverting Ornstein-Uhlenbeck process (Hansen, 1997; Butler and King, 2004; Beaulieu et al., 2012; Mahler et al., 2013; Khabbazian et al., 2016; Bastide et al., 2018). Application of these methods to empirical data has yielded mixed results; true early bursts appear to be rare (Harmon et al., 2010), though challenging to detect (Slater and Pennell, 2014) and not entirely unheard of (Slater et al., 2010; Astudillo-Clavijo et al., 2015; Slater and Friscia, 2019; Stanchak et al., 2019), while support for diversification on a rugged adaptive landscape has been recovered in some clades (e.g., Mahler et al., 2013; Benson et al., 2018; Godoy et al., 2019; Mongiardino Koch, 2021) but not in others (Law et al., 2019).

Conceptual models of adaptive radiation developed by Osborn (1902) and Simpson (1953) focused on the ecological roles that organisms play in their environments and used phenotypes as proxies, with differences in trait values between lineages being interpreted as evidence for meaningful niche differentiation (Givnish, 1997, 2015; Arnold et al., 2001). Similarly, mathematical models of clade dynamics show that traits that are directly associated with resource use evolve more quickly towards their optima and maintain lower levels of variation than other traits, particularly in the face of competition (Gavrilets and Vose, 2005; Doebeli and Ispolatov, 2017). Although this work predicts that early bursts should only occur on a rugged adaptive landscape, comparative methods are data-hungry and traits associated with resource use are labor-intensive to collect. This has led many authors (e.g., Burbrink and Pyron, 2010; Harmon et al., 2010; Slater et al., 2010; Venditti et al., 2011; Burbrink et al., 2012; Godoy et al., 2019; Benson et al., 2018) to use readily available body size data as a proxy for ecology, due to the presumed correlation between size and many aspects of life history (Peters, 1986). The correlation between body size and axes of ecological diversification is weak in many clades, though, and reliance on it for more general macroevolutionary analyses has been noted to generate misleading results (Jablonski, 1996; Slater, 2015).

Further complications arise as different models of adaptive radiation predict different relationships between evolutionary dynamics along distinct axes of resource use. Osborn’s (1902) original formulation of adaptive radiation (Osborn, 1902) was based on the idea of rapid, adaptive evolution across multiple distinct trait complexes within a narrow temporal window (Figure 1 A–C). This “simultaneous radiation” model is often associated with adaptive radiations that result from colonization of a new area or the extinction of competitors, both of which dramatically expand the range of ecological opportunities available to the radiating clade (Valentine, 1980; Walker and Valentine, 1984; Erwin, 1992). Support for time homogeneous evolutionary rates or a flat adaptive landscape associated with one axis of resource use would certainly provide evidence against simultaneous radiation, but it is also possible that adaptive radiations might instead unfold sequentially, or in stages, along distinct axes of resource use (1D–F; Diamond, 1986; Schluter, 2000; Streelman and Danley, 2003; Gavrilets and Vose, 2005; Silvertown et al., 2006b; Glor, 2010). Unfortunately, few studies have simultaneously investigated tempo and mode in phenotypic evolution across distinct ecological axes, and those to have done so have yielded contradictory results regarding the order of niche divergence (Richman and Price, 1992; Ackerly et al., 2006; Silvertown et al., 2006a; Ingram, 2011; Sallan and Friedman, 2012; López-Feráandez et al., 2013).

**Figure 1:**
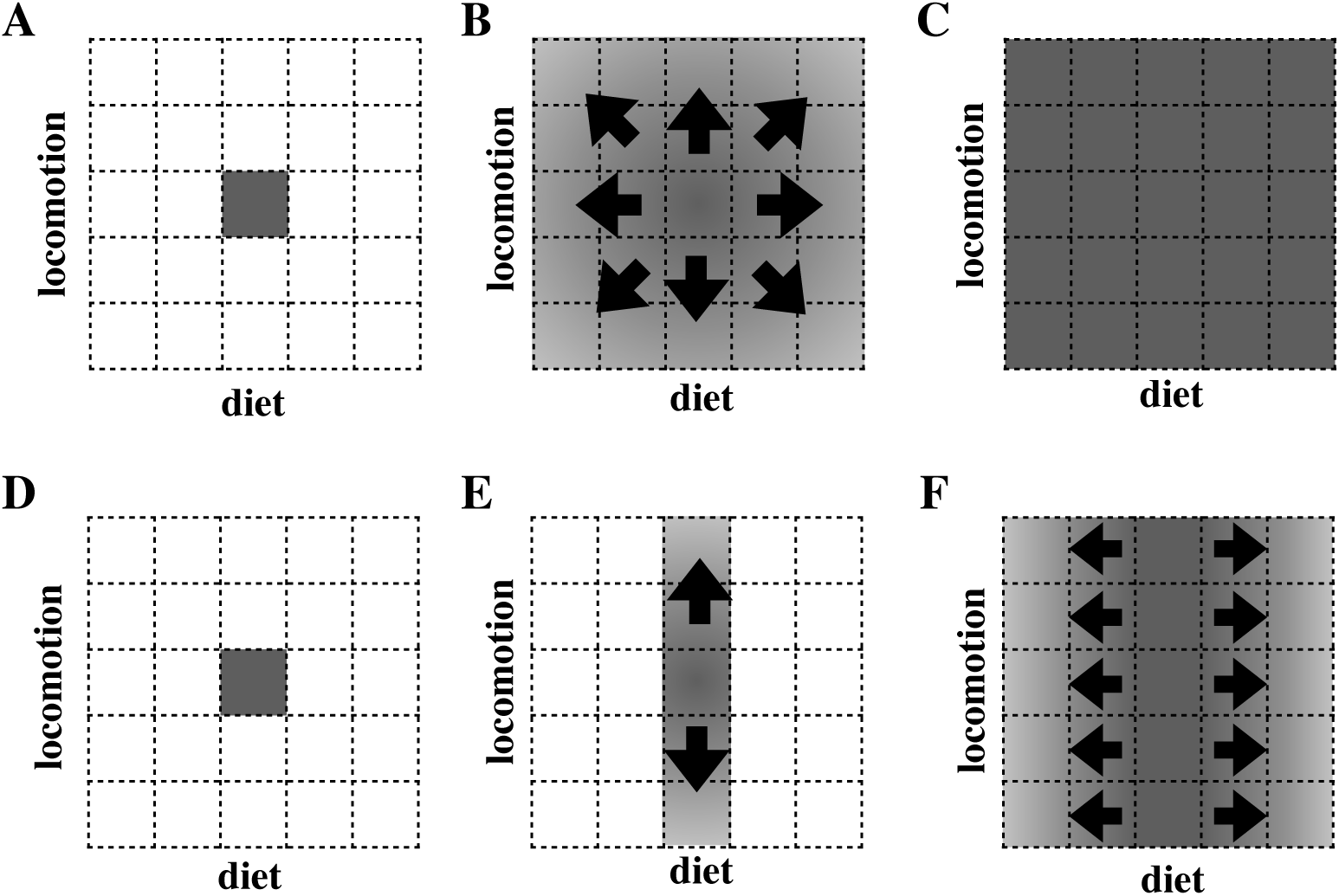
Some theories of adaptive radiation posit that a generalized ancestral form (A) should diversify simultaneously along distinct axes of resource use (here, diet and locomotor mode) (B), establishing complete functional diversity early in clade history with little subsequent change (C). Models of staged adaptive radiation differ from the simultaneous model in suggesting that the same generalized ancestor (D) should first diversify along one axis (frequently substrate-use) (E) and subsequently along another (e.g. resource-use) (F), yielding a sequential pattern of diversification in morphological and ecological data.

In this paper, I provide quantitative phylogenetic tests of the simultaneous and staged models of adaptive radiation using extant terrestrial members of the mammalian order Carnivora (dogs, cats, bears, weasels, etc.). Carnivorans are ideally suited for such a test as they are numerically diverse (*>* 200 extant species), possess a well-resolved phylogeny, and are ecologically disparate. Carnivorans span a range of dietary specializations, from plant and fruit specialists to obligate faunivores, and also exhibit a diversity of locomotor strategies, from semi-aquatic to arboreal, all of which are reflected in dental and skeletal morphology (Van Valkenburgh, 1987, 1988, 1991; Sacco and Van Valkenburgh, 2004; Friscia et al., 2007; Van Valkenburgh, 2007; Meachen-Samuels and Van Valkenburgh, 2009; Samuels et al., 2013). In an analysis of tempo and mode in mandibulodental evolution, Slater and Friscia (2019) found strong support for an early phylogenetic partitioning of dental ecomorphological variation in carnivorans, which they interpreted as support for an adaptive radiation along an axes of dietary resource use (see also Meloro and Raia, 2010). This result provides a firm basis for asking similar questions about the evolution of locomotor diversity in the clade, as well as the relationship between macroevolutionary dynamics and the topography of the underlying adaptive landscapes for these traits. Here, I specifically attempt to address three key questions related to the nature of adaptive radiations and our ability to detect them from comparative data. First, I ask whether carnivoran postcranial functional traits exhibit a signature of early, rapid morphological diversification consistent with traditional models of early-burst adaptive radiation and the signal present in some mandibulodental metrics. Then, I ask whether traits associated with dietary resource use, locomotor mode, and body size diversify simultaneously or sequentially and, if the latter, in which order do carnivorans diversify. Finally, I quantify the topography of the underlying macroevolutionary adaptive landscapes associated with these traits to ask whether variation in the tempo of trait diversification is associated with presence or absence of distinct adaptive peaks. My results indicate that diet, but not locomotor mode or body mass, was the critical axis of resource use exploited by the radiation of crown-group carnivorans.

## Materials and methods

### Phylogenetic Framework

A well-supported, time-scaled phylogenetic framework is essential for testing hypotheses regarding the tempo and mode of morphological diversification. For this study, I used the maximum clade credibility tree for extant and a few recently extinct terrestrial carnivorans generated by Slater and Friscia (2019). The tree topology and its branch lengths were jointly inferred using BEAST v.2.3.2 (Bouckaert et al., 2014) from a supermatrix of 30 nuclear loci and protein-coding mitochondrial genes for 235 taxa under a relaxed uncorrelated log-normal clock (Drummond et al., 2006). The unresolved fossilized birth-death process (Heath et al., 2014; Gavryushkina et al., 2014, 2016) was used as a tree prior, and 250 phylogenetically constrained fossil taxa with associated stratigraphic ages were used to calibrate the relaxed molecular clock. The resulting topology is well-supported (94% of splits have posterior probabilities *>* 0.9) and 95% highest posterior density intervals for node ages are narrow (mean = 2.65 Myr).

### Ecomorphological Data

Species mean values for 28 mandibulodental linear measurements as well as 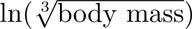 body mass) for 198 species of extant carnivoran were taken from Slater and Friscia (2019). These were supplemented with 24 new measurements of limbs, scapulae, and pelves from 547 specimens spanning 135 species of extant carnivorans (mean = 4, range = 1-15 specimens per species) housed in the mammalogy collections of the National Museum of Natural History (Smithsonian Institution), Washington D.C., and the Field Museum, Chicago. Measurements were taken using Mitutoyo digital calipers to 0.01mm precision for measurements *<* 250mm, or with a tape measure to 1mm precision for measurements *>* 250mm and were selected to allow the computation of 16 ratio-based functional indices (Table 1) that have previously been identified as useful in distinguishing carnivores with distinct locomotor strategies (Davis, 1964; Van Valkenburgh, 1987; Samuels et al., 2013; Gould, 2014). Measurements of the metacarpals and tarsals, though highly informative in distinguishing among mammals with distinct locomotor modes (Samuels et al., 2013; Nations et al., 2019), were not taken here as these elements were infrequently available in osteological preparations of small-bodied taxa and their inclusion would therefore have reduced the number of species that could be sampled for macroevolutionary analyses. I corroborated that postcranial measurements allowed the discrimination of carnivorans with different substrate use preferences by performing a linear discriminant analysis using the lda function from the MASS library (Venables and Ripley, 2002, Supplementary File A). Each species with available post-cranial measurements was assigned to one of Eisenberg’s (1981) substrate use categories (Terrestrial, Scansorial, Arboreal, Natatorial, Semi-Fossorial) following Polly (2010), with some minor edits based on literature review. Leave-one-out cross-validation (Supplementary File A) showed that most taxa can be assigned to their locomotor grouping with relatively high posterior probability, and that where mis-classifications occur, they tended to place taxa into similar locomotor groupings (e.g., scansorial taxa classified as terrestrial).

**Table 1:**
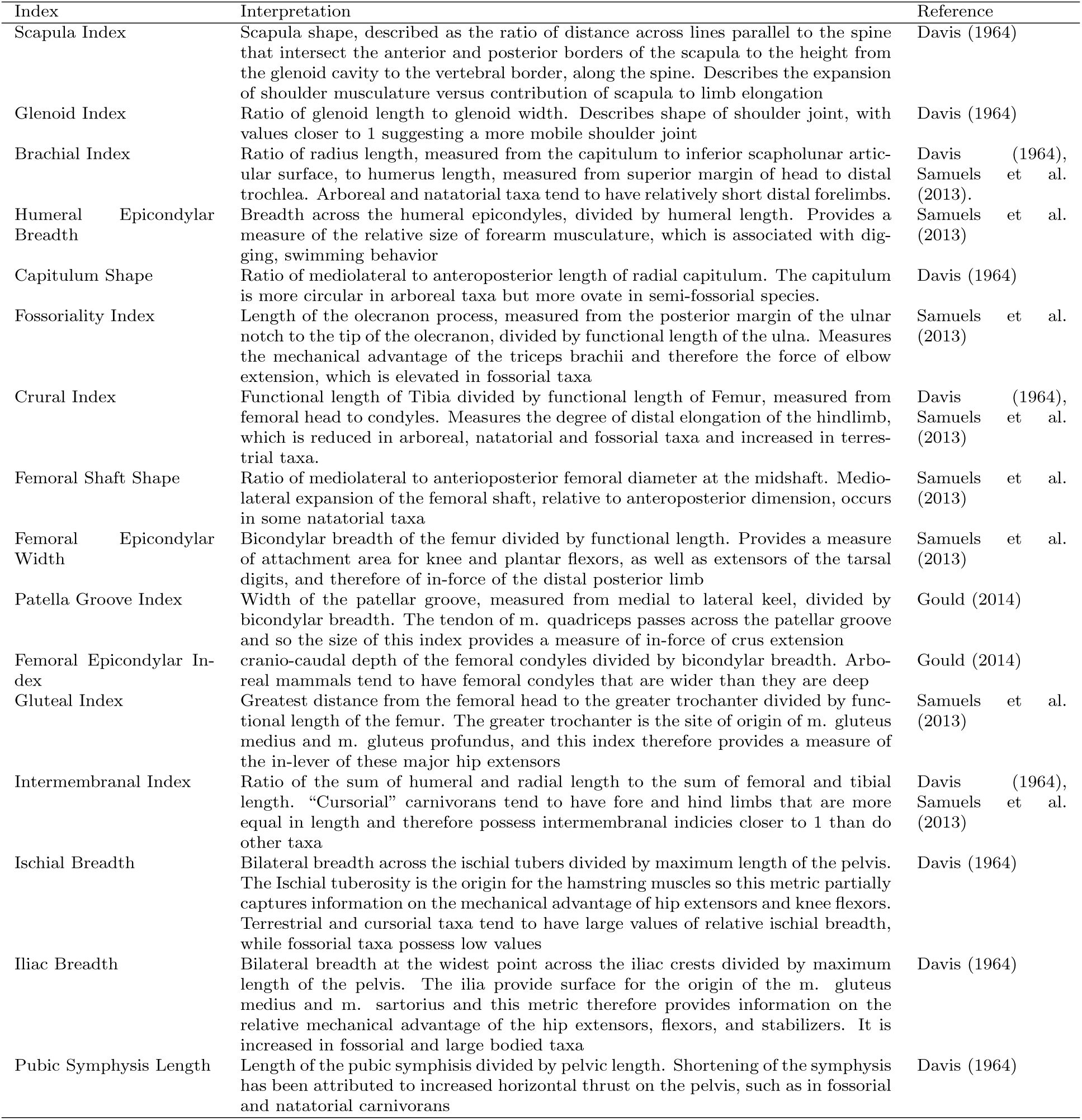
Post-cranial functional indices computed for carnivoran species in this study, along with their constituent measurement, functional interpretation, and source(s)

### Macroevolutionary models of post-cranial evolution

Slater and Friscia (2019) recovered strong support for an “early burst” of evolution in carnivoran molar traits but not in other mandibulodental characteristics or body mass. They considered this result to be consistent with adaptive radiation along dietary axes and cautioned that different functional indices may provide different macroevolutionary signals. To assess whether locomotor diversification in carnivorans occurred along one or a few axes of functional post-cranial variation, I evaluated the fit of the same three macroevolutionary models (constant-rates Brownian motion, single-stationary-peak Ornstein-Uhlenbeck model, and early burst) to each of the 16 functional trait indices described above by using the fitContinuous() function from the geiger library (Pennell et al. 2014, Supplementary File B). To account for measurement error I added sampling variances to the diagonals of the model-specific variance-covariance matrices. Relative model fit was assessed by computing small-sample corrected Akaike Weights, *w_A_*, for each model.

### The Accumulation of Ecomorphological Disparity Through Time

The simultaneous radiation model predicts that diversification along dietary and substrate-use axes should occur at approximately the same time, while the staged radiation model predicts that locomotor evolution should precede dietary diversification. To differentiate between these two forms of adaptive radiation, it is therefore necessary to assess which axis of trait variation, if any, was exploited first. In the paleobiological literature, weighted average times, often referred to as the Center of Gravity (CG), have long been used as measures of bottom- or top-heaviness in disparity and diversity profiles (Gould et al., 1977; Foote, 1991). Past levels of disparity cannot be inferred from phylogenies of extant taxa, but the partitioning of variation over phylogeny can be evaluated using Disparity Through Time analysis (DTT; Harmon et al., 2003), and this method lends itself well to CG descriptors.

DTT analyses track the standardized average morphological variance of subclades that are extant at time points (=node ages) spanning the root of a phylogeny to the final divergence event in the tree, where the standardization is relative to the total variance of the clade (Figure 2). If *d_i_* is the standardized average subclade disparity at time *t_i_*, then the center of gravity of the DTT profile can be computed as

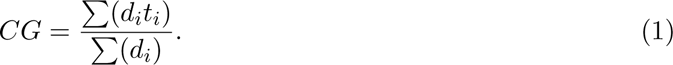

**Figure 2:**
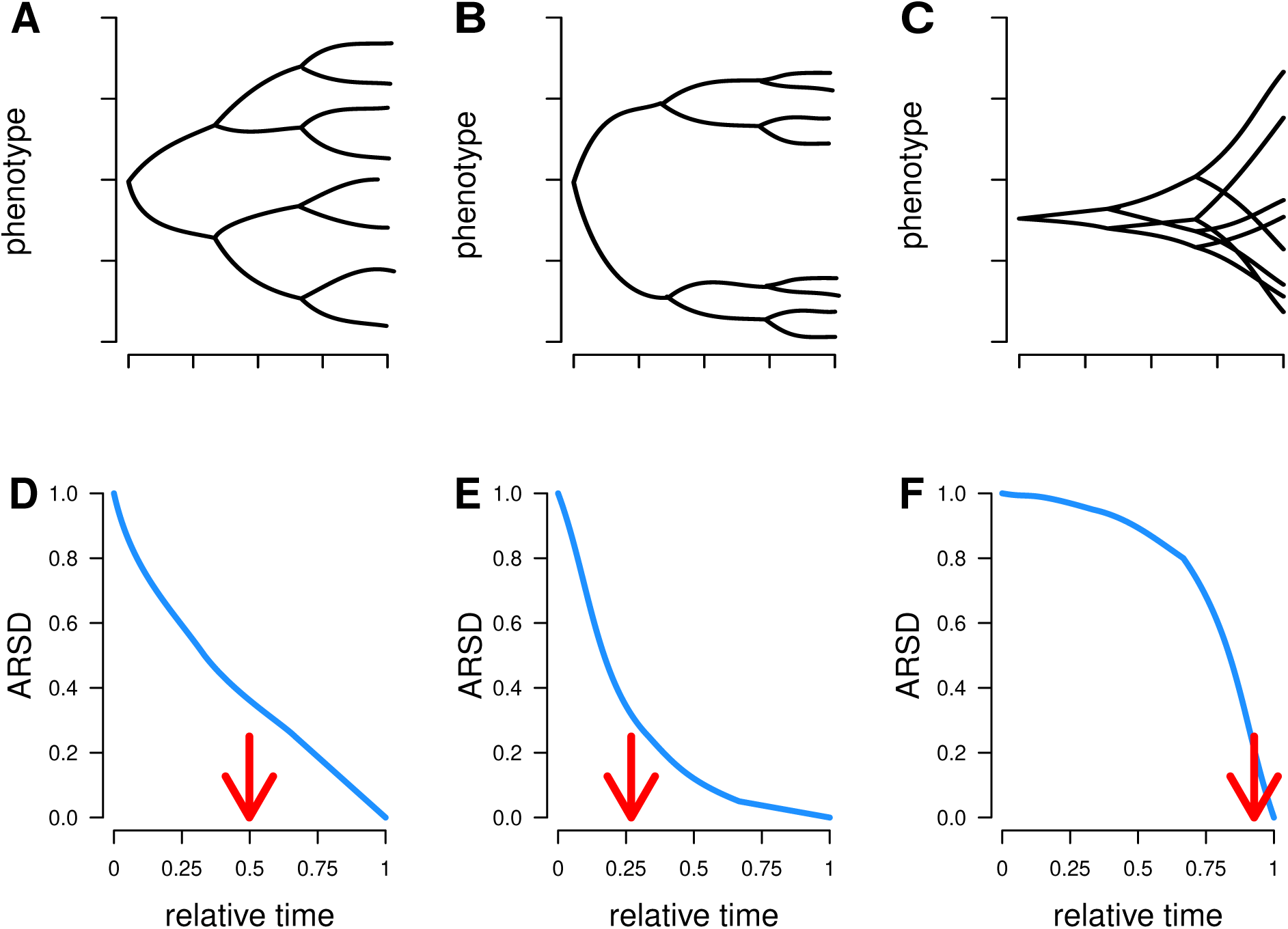
When the mode of phenotypic evolution is consistent with a random walk, phenotypic variation accumulates gradually over phylogeny (A), leading to a pattern in which the average relative subclade disparity through time (ARSD) declines steadily from 1 (i.e. all variance within a single clade) to zero (all clades are single tips) as time progresses from root to tip (D) and a center of gravity (red arrow) that falls at approximately the midpoint. Under an “early burst” scenario, trait variation does not accumulate steadily but rather, seems to accumulate early in the history of the clade (B), yielding an ARSD curve that drops precipitously and then levels out (E), and a center of gravity that falls below the midpoint of clade history. Evolutionary modes such as “late bursts” or time constant rates in a bounded space (e.g., constraints) yield a pattern in which phenotypic variation accumulates towards the tips of the tree (C) and an ARSD curve that declines very slowly until late in clade history (F), with a very high center of gravity.

Because the average subclade disparity is standardized relative to total clade variance, it declines from a value of *d_t_* = 1 at relative time *t* = 0 (i.e. along the root edge of the phylogeny) to *d_t_* = 0 at the final branching event on the tree where each extant lineage is an individual species. This conveniently yields an expectation that CG should fall at *t ≈* 0.5 (see Supplementary File D for a simulation-based confirmation), with low CGs representing an early partitioning of variation among subclades and high CGs representing the late maintenance of high within subclade variation. Of course, the exact expected CG for any clade will depend on the shape of the tree (Foote, 1993) and this expectation can be conveniently assessed and compared with the empirical CG via simulation under a constant rates process. This approach is conceptually similar to the Divergence Order Test (DOT) of Ackerly et al. (2006), which calculates the center of gravity for unstandardized independent contrasts (Felsenstein, 1985) to determine whether large divergences in trait values occur early or late in clade history. A potential limitation of the DOT is that contrasts are computed using ancestral state estimates that are, themselves, inferred under a time-homogeneous model of evolution (Sallan and Friedman, 2012). Furthermore, the DOT explicitly assumes that adaptive radiation results from decelerating rates of phenotypic evolution, whereas similar patterns may result from myriad other process, including evolution on a rugged adaptive landscape (e.g., Revell et al., 2005; Harmon et al., 2021). Unlike the DOT, the DTT center of gravity does not explicitly assume an underlying evolutionary process but, instead, provides a means of describing and statistically evaluating whether the temporal structure of morphological variation within and among clades is consistent with the hypothesis of early variance partitioning among clades (Foote, 1996).

I calculated CGs for mandibulodental and post-cranial data, as well as for body mass, and evaluated their deviation from constant-rates expectations using 9999 datasets of equivalent dimensionality simulated under a constant-rates Brownian motion process (Supplementary File C). I used the original linear measurements, rather than the functional ratios, for the manidibulodental and post-cranial datasets. However, in order to remove consideration of overall size and to focus on ecomorphological shape, these linear dimensions were transformed into log-shape variables (Mosimann, 1970; Darroch and Mosimann, 1985) by dividing each by the geometric mean measurement for each species and then taking the natural log of this ratio. The high dimensionality of the mandibulodental and post-cranial traits poses challenges for efficient and accurate macroevolutionary inference. To reduce data dimensionality, I therefore performed a principal components (PC) analysis on the correlation matrices for each set of log-shape variables using the prcomp function in the stats library and used the broken-stick method (Frontier, 1976) to determine the optimal number of PCs to retain for further analysis. This approach tends to identify a smaller number of axes than other methods, such as the Kaiser-Guttman criterion (axes with eigenvalues *≥* 1) or minimum/cumulative values of relative eigenvalues, but outperforms these approaches in simulation tests (Jackson, 1993). CGs were finally compared across sets of traits to determine the relative order of divergence.

### The Adaptive Landscape of Carnivoran Ecomorphological Diversificaton

Simple macroevolutionary models and centers of gravity provide insights into the phylogenetic partitioning of morphological variation and hint at the kinds of evolutionary processes that might be responsible, but they do not allow us to discriminate between modes of adaptive radiation based purely on declining rates and those resulting from adaptive evolution on a rugged landscape. To understand the context for carnivoran ecomorphological evolution, I therefore estimated the macroevolutionary landscapes for mandibulodental and post-cranial traits, as well as body mass, using the phylogenetic expectation-maximization approach implemented in the PhylogeneticEM library (Bastide et al., 2018). This approach assumes that trait evolution follows an Ornstein-Uhlenbeck process in which traits evolve towards a primary optimum or, in the case of multivariate trait evolution, a vector of primary optima (*θ*). The method then attempts to identify the presence and location of shifts in the vector of primary optima. Univariate traits evolve according to a standard Ornstein-Uhlenbeck process (Hansen and Martins, 1996b; Hansen, 1997) but traits can also be modeled in a true multivariate framework, with variances and covariances specified by an evolutionary rate matrix **R**, though a single strength of attraction, *α*, must then be shared by all traits.

As with DTT analyses, I used PC axes retained by the broken stick criterion for each multivariate dataset. The maximum number of shifts allowed, K max, was set to 30, based on preliminary trials; the nbr *α* argument, which determines the resolution of the grid of *α* values evaluated, was set to 100; the LINselect option was used for model selection. For convenience of interpretation, *α* values were converted into phylogenetic half-lives 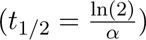, which correspond to the time taken for a trait to move halfway from its ancestral value to a new primary optimum after a regime shift and the diffusion parameters of the stochastic component of the OU process were converted to vectors of equilibrium variances 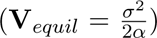, which represent the expected variances of the process along each axis of shape variation at stationarity (Hansen, 1997).

## Results

### Macroevolutionary model fitting

Contrary to predictions of both the simultaneous and staged models of adaptive radiation, I found no evidence for early bursts of locomotor adaptation within Carnivora. Brownian motion or SSP models were uniformly recovered as best-fitting across the 16 functional indices, with support for the early burst model never exceeding that of BM or SSP (Table 2), and maximum likelihood estimates of the early burst exponential rate decline parameter were never large (maximum *r* =-0.0016, equivalent to a rate half-life of 433myr). Maximum likelihood estimates of the *α* parameter of the OU process are relatively small, at most corresponding to a phylogenetic half-life of 6.3 myr (Femoral Shape), and more typically in the range of 23-34myr.

**Table 2:** Akaike Weights for the three models (BM: Brownian motion; SSP: single stationary peak; EB: early burst) fitted to each Functional Index. The weight corresponding to the best fitting model for each index is bolded. Parameter estimates are provided for the best fitting model; note that the *α* parameter is only relevent to SSP models.

| Functional Index | BM | SSP | EB | Root | Rate | $\alpha$ |
| --- | --- | --- | --- | --- | --- | --- |
| Scapula Index | **0.59** | 0.21 | 0.21 | 0.85 | $2.9*10^{-4}$ | - |
| Glenoid Shape | 0.01 | **0.99** | 0.00 | 0.69 | $2.1*10^{-4}$ | 0.05 |
| Brachial Index | 0.12 | **0.83** | 0.04 | **0.84** | $6.3*10^{-4}$ | 0.03 |
| Humeral Epicondylar Breadth | **0.52** | 0.29 | 0.18 | 0.25 | $9.6*10^{-5}$ | - |
| Capitulum Shape | 0.38 | **0.49** | 0.13 | 1.46 | $4.9*10^{-4}$ | 0.03 |
| Fossoriality Index | 0.16 | **0.78** | 0.06 | 0.14 | $7.1*10^{-5}$ | 0.03 |
| Crural Index | 0.37 | **0.50** | 0.13 | 0.93 | $5.2*10^{-4}$ | 0.02 |
| Femoral Shaft Shape | 0.00 | **1.00** | 0.00 | 1.12 | $1.0*10^{-3}$ | 0.11 |
| Femoral Epicondylar.width | **0.52** | 0.30 | 0.18 | 0.19 | $3.0*10^{-5}$ | - |
| Patella Groove Index | **0.44** | 0.4.0 | 0.16 | 0.43 | $6.6*10^{-5}$ | - |
| Femoral Epicondylar Index | 0.23 | **0.69** | 0.08 | 0.01 | $1.0*10^{-6}$ | 0.03 |
| Gluteal Index | **0.44** | 0.4.0 | 0.16 | 0.18 | $2.9*10^{-5}$ | - |
| Intermembranal Index | 0.39 | **0.47** | 0.14 | 0.84 | $2.0*10^{-4}$ | 0.02 |
| Ischial Breadth | 0.12 | **0.83** | 0.04 | 0.58 | $4.6*10^{-4}$ | 0.03 |
| Iliac Breadth | **0.58** | 0.21 | 0.21 | 0.64 | $6.0*10^{-4}$ | - |
| Pubic Symphysis Length | **0.57** | 0.23 | 0.2.0 | 0.27 | $1.5*10^{-4}$ | - |

### The Accumulation of Ecomorphological Disparity Through Time

Principal components analysis of the mandibulodental log-shape variables yielded 2 significant PCs according the broken stick analysis. A plot (Fig. 3A) of the shape space defined by PC1 (*∼*64% of the variance) and PC2 (12.5% of the variance) shows a generally obvious phylogenetic partitioning, with more carnivorous clades (e.g., Felidae, Hyaenidae) occupying negative PC1 positions due to a combination of deep and stout jaws, large jaw muscle moment arms, robust canines and small or absent second and third molars, the bears (Ursidae) occupying positive PC1 and negative PC2 positions due to their large posterior lower molars, and weasels (Mustelidae) and skunks (Mephitidae) occupying positive PC1 and positive PC2 positions due to the large talonid of the lower first molar. Mongooses (Herpestidae), civets (Viverridae), and dogs (Canidae) fall closer to the origin of this plot due to their relative plesiomorphic dental conditions. PC loadings for retained PCs are provided in Supplementary Table C1.

**Figure 3:**
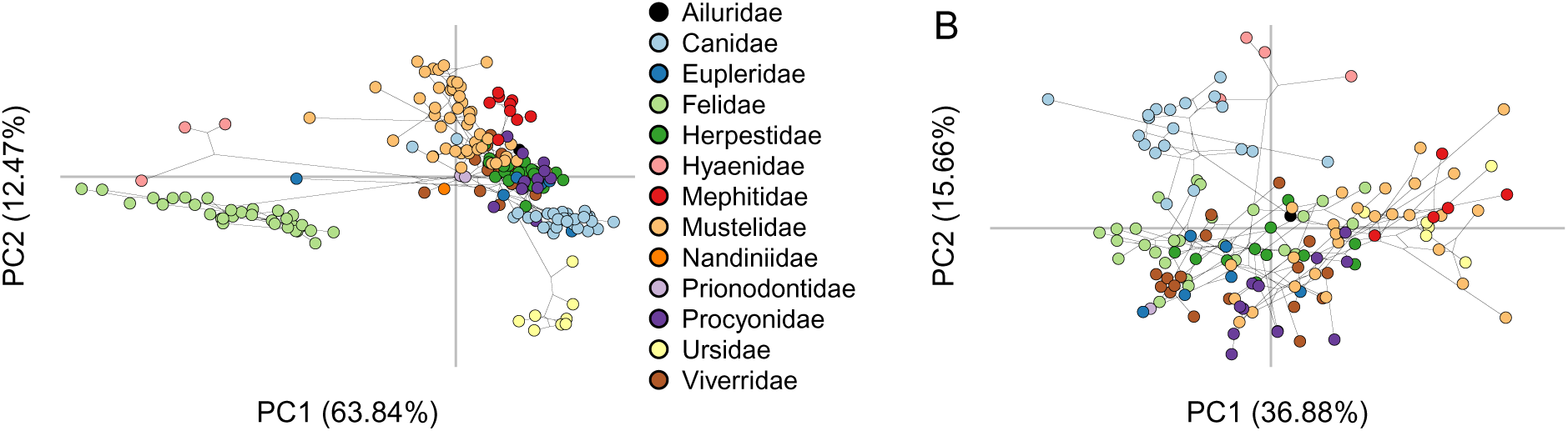
Principal components analysis of log shape variables from the mandibulodental log shape variables (A) shows a strong phylogenetic partitioning. The shape space defined by PCs 1 and 2 of log-shape variables from the post-cranial skeleton (B) shows slight separation of some clades and functional groups along PCs 1 and 2, but with much more overlap than is found for the mandibulodental data.

Three principal components were retained from analysis of the post-cranial log-shape variables. A plot of PC1 and PC2 (Fig. 3B) shows some phylogenetic partitioning, with Canidae and Hyaenidae in particular clustering in the positive PC2 region due to their especially tall and narrow scapulae, pelves with broad ischia and a short symphysis, and long distal limb elements. A functional signal is also clearly present, with more arboreal taxa from several clades (*Bassaricyon* sp., *Arctogalidia*, *Martes*, *Cryptoprocta*) tending towards negative PC2 scores due to their short broad scapulae, narrow pelves with a long symphysis, and broad distal femora. More terrestrial (e.g. *Ursus* sp.), semi-fossorial (e.g. *Taxidea*, *Meles*, Mephitidae), and natatorial (otters; Lutrinae) species trend towards the positive end of PC1 due to their short broad limbs and robust girdles, while more scansorial taxa (Felidae, some Viverridae and Eupleridae) trend towards negative PC1 scores due to more elongate and gracile limbs. PC loadings for retained PCs are provided in Supplementary Table C2 .

The average subclade disparity through time profile for mandibulodental traits is broadly consistent with constant-rates expectation for the first 15 million years of carnivoran history, as indicated by a curve (solid line) that tracks the mean of the Brownian motion simulations (dashed line; Figure 4A). However, mandibulodental subclade disparity shows a substantial drop in the mid-Oligocene and lies at the lower bound of the constant-rates expectation (shaded area) for the rest of the Cenozoic (Figure 4A). The Center of Gravity for mandibulodental average subclade disparity appears slightly top-heavy (CG=0.58); however, the expected CG is dependent on the structure of the phylogeny (Supplementary File D) and is, in fact, significantly lower than expected under a constant rates process (95% quantiles under BM = 0.60-0.72 ; *P* =0.008;Fig. 4D). As suggested by the disparity through time curve (Figure 4B) the Center of Gravity for post-cranial traits (Fig 4C) is much higher than that for mandibulodental traits and is significantly higher than expected under a constant-rates process (CG = 0.68, *P* = 0.002; Figure 4E). The partitioning of body mass variation among and within clades is entirely consistent with a constant rates process (Fig 4C), with a CG that is virtually indistinguishable from the BM expectation for a univariate trait (CG = 0.7, Expectation = 0.7, *P* = 0.177; Figure 4F). The order of divergence, from earliest to most recent, therefore appears to be mandibulodental traits, body mass, and then post-cranial traits.

**Figure 4:**
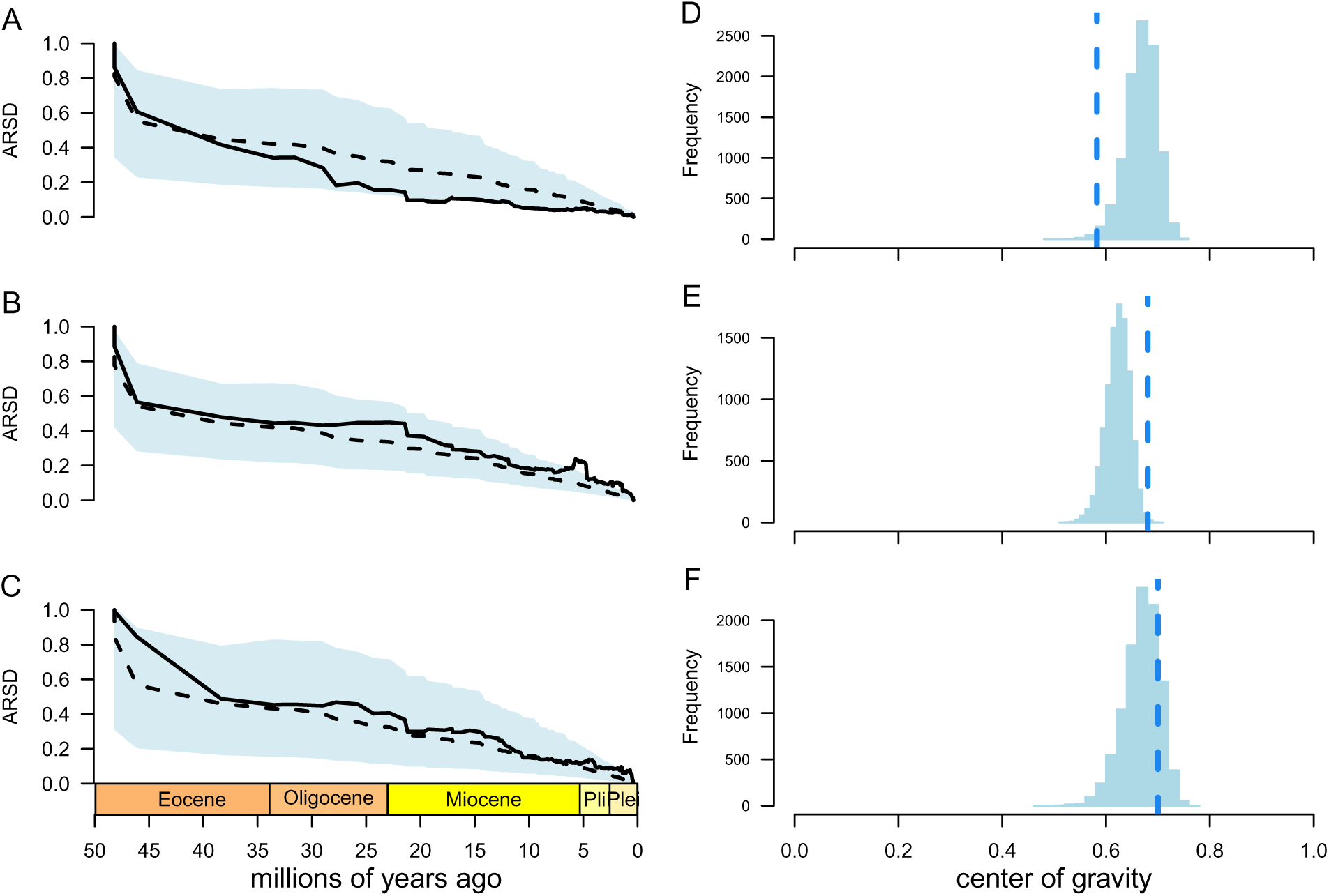
Disparity through time plots for carnivoran mandibulodental (A), post-cranial (B) and body mass (C) data. The solid line is the average relative sub-clade disparity for the data set, the dashed line is the median from 9999 datasets simulated under a constant rates process, and the shaded area corresponds to the 95% quantiles of the simulated data. Shown on the right (D-F) are the corresponding centers of gravity for the simulated datasets (solid bars) and the center of gravity for the trait data (dashed vertical line)

### The Adaptive Landscape of Carnivoran Ecomorphological Diversificaton

The dietary adaptive landscape estimated from the first two mandibulodental PC axes by the PhylogeneticEM algorithm contains 21 peaks that are distinct from the ancestral regime. Note that PhylogeneticEM does not identify and collapse convergent shifts (shifts to similar optimal morphologies) and so some of these peaks may overlap one another (see below). The optimal configuration is shown in Figure 5A; 72 equivalent configurations were recovered that differ in the local arrangement of shifts (Supplementary File C). For example, shifts may be swapped between sister taxa (sequential vs independently acquired). The strength of the pull to these optima is relatively strong (*α* = 0.34), corresponding to a phylogenetic half-life of *∼ t*_1/2_ = 2.04 myr. Combined with estimates for the stochastic diffusion component of the OU process, this yields very small equilibrium variances along each axis of shape variation (**V***_equil_* = [0.78, 0.20]).

**Figure 5:**
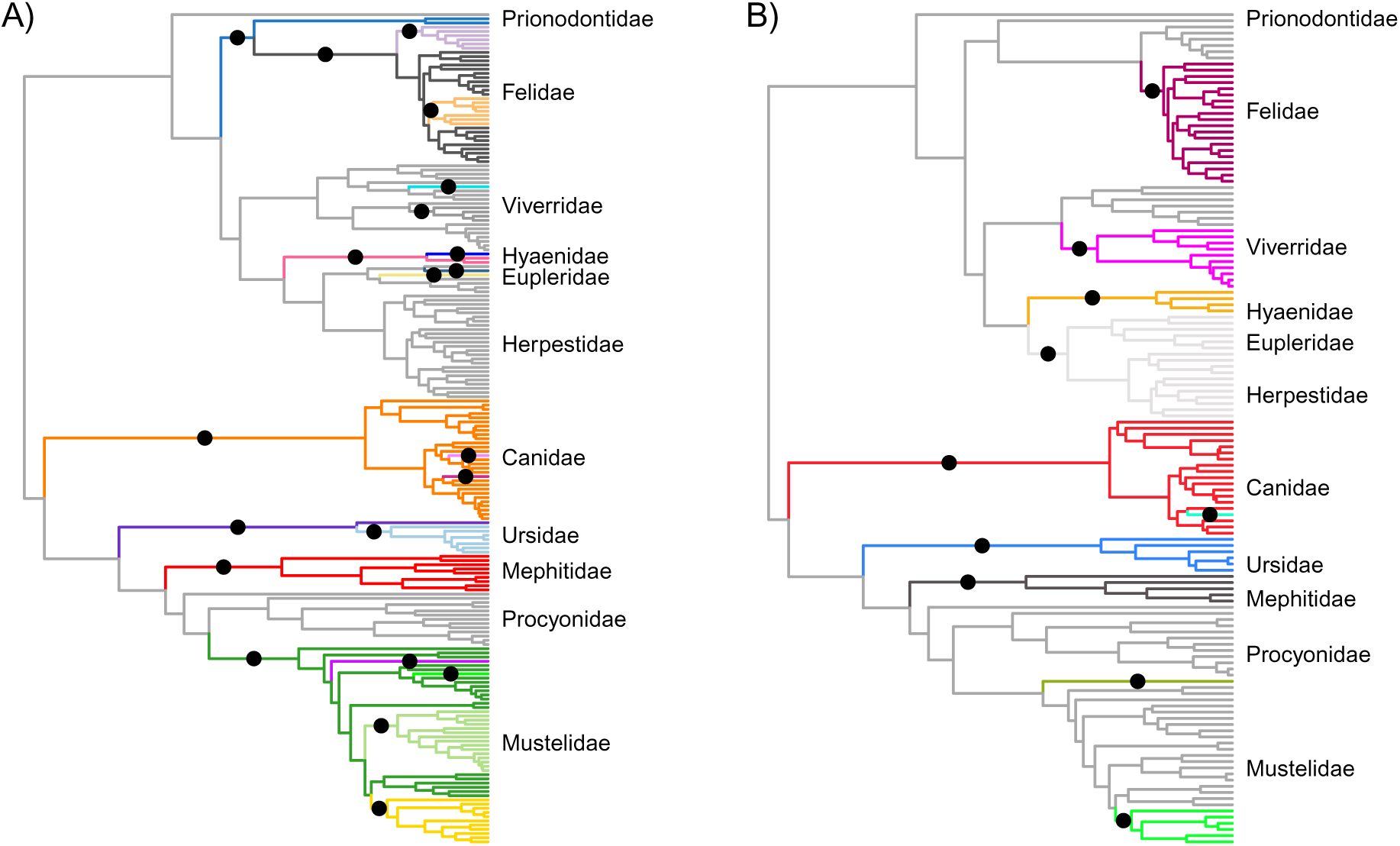
Carnivoran phylogeny showing the 25 adaptive peak shifts for mandibulodental traits identified by the phyloEM approach. Black circles are placed at the midpoint of each branch along which a peak shift is inferred to have occurred, with branches inheriting and retaining that peak colored distinctly. Clade names are provided to orient the reader to shifts described in the results and are not intended to be exhaustive.

Within feliform carnivorans, peak shifts are recovered along the branches leading to an unnamed clade comprising Prionodontidae + Felidae, Felidae, Pantherinae, the Lynx and Puma clades within Felinae, Hyaenidae, and terminal branches leading to the spotted hyaena *Crocuta crocuta* and the Malagasy Fossa *Cryptoprocta ferox*. All of these taxa are hypercarnivorous relative to other members of the suborder. Additional shifts are found along the branch leading to the binturong (*Arctictis*), the Viverrinae (exclusive of *Vivericula indica*) and *Eupleres goudotii*, representing dietary shifts towards frugivory and insectivory. Among caniform carnivores, shifts are recovered along the branch leading to Canidae and within the family on terminal branches corresponding to the hypercarnivores *Speothos venaticus* and *Cuon alpinus*, Ursidae, Tremarctinae + Ursinae (i.e., Ursidae minus the giant panda *Ailuropoda*), Musteloidea, Mephitidae, Mustelidae, the honey badger *Mellivora capensis* and the wolverine *Gulo gulo*, Mustelinae and Lutrinae. These shifts represent a more diverse set of dietary ecologies, ranging from more carnivorous (Mustelinae, *Gulo*) or piscivorous (Lutrinae) to more omnivorous (Ursidae).

The adaptive landscape for post-cranial traits showed fewer peaks than that for mandibulodental traits, with 10 shifts identified in total, and 3 equivalent configurations. For the optimal configuration (Figure 5B), shifts occur within Feliformia along the branch leading to Felinae, Viverrinae, Hyaenidae, and Eupleridae + Herpestidae. With the exception of Hyaenidae, which is more terrestrial and “cursorial” and, perhaps, Prionodontidae + Felidae, which is more scansorial, a functional signal here is not apparent. Within Caniformia shifts appear to have more obvious functional interpretation. Shifts are identified along the branches leading to the terrestrial Canidae (with an additional shift along the branch leading to the long-legged maned wolf *Chrysocyon brachyurus* ), the large-bodied, plantigrade Ursidae, the semi-fossorial skunks (Mephitidae) and American badger (*Taxidea taxus*), and the semi-aquatic Lutrinae. In addition to containing fewer peaks, the post-cranial adaptive landscape is less rugged than that for mandibulodental traits. The rate of adaptation (*α* = 0.161) translates to a phylogenetic half-life of *t*_1/2_ = 4.3 myr. This also results in equilibrium variances that are much larger that those for the dental traits (**V***_equil_* = [2.29, 1.22, 1.09]).

A flat adaptive landscape emerges for body mass, with no shifts detected and a strength of pull the mean that is an order of magnitude smaller than that for dental and post-cranial traits (*α* = 0.0177*, t*_1*/*2_ = 39 myr).

## Discussion

Models of adaptive radiation were originally developed to explain the early, rapid appearance of fundamentally distinct (“essentially discontinuous”; Simpson, 1953, p.223) modes of life within diversifying clades in the fossil record (Osborn, 1902). A modern focus on the ”early rapid” component of these models has led to the development of numerous methods for assessing temporal variation in macroevolutionary rates (Blomberg et al., 2003; Harmon et al., 2003; Freckleton and Jetz, 2009; Harmon et al., 2010; Venditti et al., 2011) but yielded limited support for this mode of diversification in body size and shape data. The concept of an underlying adaptive landscape that links form to fitness, while also crucial to these models, has received less attention in modern phylogenetic tests of the early burst adaptive radiation hypothesis (but see Mahler et al., 2013; Aristide et al., 2018; Mongiardino Koch, 2021), and it therefore remains unclear whether adaptive radiations unfold simultaneously across distinct axes of resource use, or sequentially with some axes being exploited before others. Using methods that focus on the temporal accumulation of morphological variation (Harmon et al., 2003; Slater and Pennell, 2014) and the topography of the underlying adaptive landscape (Bastide et al., 2018), rather than variation in rates, I found that significant early partitioning of mandibulodental morphological variation across carnivoran phylogeny occurs on a rugged adaptive landscape with multiple peaks, consistent with classic ideas about adaptive radiation (Osborn, 1902; Simpson, 1953; Valentine, 1969). Although strong support for this mode of adaptive radiation is present in traits related to diet, its signal is not present in body mass data or for traits related to locomotor behavior and substrate use. These findings suggest that the nature of adaptive radiation is more complex that simple univariate tests might suggest, and provide some hints regarding fruitful directions that future work might explore.

### Carnivorans Adaptively Radiated on a Rugged, Multi-Peak Dietary Landscape

Competition for dietary resources has long been viewed as a critical factor promoting the evolution of morphological innovations in some of the most iconic examples of adaptive radiation (Darwin, 1845; Lack, 1947; Grant, 1981; Schluter and Grant, 1984; Osborn, 1902). Theoretical models lend support to this hypothesis by predicting that early rapid evolution along axes of resource use should promote speciation and stable coexistence (Gavrilets and Vose, 2005; Doebeli and Ispolatov, 2017), while genomic evidence has started to accumulate that indicates that relatively simple, and sometimes repeatable, genetic or transcriptomic “key innovations” can produce morphological features associated with distinct dietary specializations (Berenbaum et al., 1996; Lamichhaney et al., 2015; Vizueta et al., 2019; McGee et al., 2020). Carnivorans and their extinct stem lineages (Carnivoramorpha), are characterized by the presence of a carnassial pair, a modified upper fourth premolar and lower first molar that form a scissor-like pair of blades for efficiently cutting tough materials such as skin and connective tissue, which first appeared in the fossil record during the latest Paleocene (Soĺe et al., 2016). Although carnassials have evolved multiple times in mammalian evolutionary history, the true innovation in Carnivoramorpha, relative to extinct mammalian clades such as Creodonta and Borhyaenidae or the extant dasyurid marsupials (Dasyuridae), is that carnassialization is restricted to a single locus and that the basin-like posterior part of the lower carnassial tooth is free to occlude with the non-carnassial upper first molar (Van Valkenburgh, 1999, 2007). By varying the proportion of the lower carnassial given to slicing (the anterior trigonid) versus grinding (the posterior talonid), carnivorans can therefore optimize efficiency at processing different types of food, ranging from vertebrate flesh to fruits or even bamboo stems (Wesley-Hunt, 2005; Van Valkenburgh, 2007). Ecomorphological (Van Valkenburgh, 1988, 1991, 2007), and behavioral (Van Valkenburgh, 1996) evidence supports a prominent role for molar morphology in determining dietary resource use, and comparative evidence for early burst-like dynamics in carnassial shape has been previously recovered (Meloro and Raia, 2010; Slater and Friscia, 2019). Recent work (Asahara et al., 2016) has also demonstrated that carnivoran-specific patterns of selection on the *Bmp7* gene are associated with correlations between the relative size of the second molar and the size of the talonid relative to the trigonid, both of which correlate with dietary ecology. This may have permitted carnivorans a greater degree of evolutionary versatility in their dentition, such as lower second molars that are larger than the lower first and third molars in ursids, than for other mammalian clades which are constrained by more fundamental developmental rules (Kavanagh et al., 2007; Polly, 2007).

As noted by Slater and Friscia (2019), support for early-burst dynamics in datasets comprising only extant taxa could potentially arise due to selective pruning of phylogeny by extinction, yielding a phylogenetically conservative distribution of traits that only the incorporation of fossil evidence could overturn (Slater et al., 2012). Although the distribution of multivariate mandibulodental trait values is phylogenetically conservative for carnivorans, the relatively fast rate of adaptation estimated by phylogeneticEM indicates that this result is not due to temporally declining rates per se but, rather, that the evolution of mandibulodental disparity in carnivorans occurred on a rugged adaptive landscape as predicted by qualitative (Osborn, 1902; Simpson, 1944, 1953) and quantitative (Gavrilets and Vose, 2005) models of adaptive radiation. The rich carnivoran fossil record could and should be leveraged to further interrogate this finding, but it is unclear whether this would result in a significant overturning of it, as selective extinction should increase, rather than decrease, phylogenetic signal in comparative data. It also remains unknown whether this mode of diversification is likely to be unique to carnivorans among mammalian orders due to their novel dental anatomy, or whether it represents a more general pattern of ecomorphological diversification (Van Valen, 1971). Fulwood et al. (2021) found no support for early burst patterns in molar topographic complexity data across Lemuriformes (Primates), but Aristide et al. (2018) found evidence for distinct adaptive peaks in cranial shape data for Platyrrhini, driven largely by variation in the craniofacial skeleton that is associated with multidimensional niche membership. Outside of Primates, much comparative macroevolutionary work has relied on body mass, which is only weakly predictive of broad dietary niche (Price and Hopkins, 2015; Grossnickle, 2020), and there remains much potential for rigorously evaluating hypotheses regarding modes of evolutionary diversification across mammalian phylogeny using ecomorphological data.

### The Locomotor Landscape is Less Rugged than the Dietary Landscape … At Least for Carnivora

Conceptual and theoretical models of staged adaptive radiation have tended to assume that divergence along habitat-use axes should precede divergence along axes of dietary resource use (Diamond, 1986; Streelman and Danley, 2003; Gavrilets and Losos, 2009). Empirical evidence in favor of this form of staged radiation is mixed (Richman and Price, 1992; Ackerly et al., 2006; Silvertown et al., 2006a; Sallan and Friedman, 2012; Ingram, 2011), but has been unclear whether this lack of support reflects real underlying evolutionary dynamics or limiting assumptions regarding rate constancy or the estimation of ancestral states (Glor, 2010). Based on the methods employed here, there is no evidence for an early partitioning of post-cranial morphological variation among carnivoran clades, despite the associated adaptive landscape exhibiting several distinct peaks. It is possible that the lack of signal for early rapid evolution recovered here reflects the omission of measurements from ecologically important post-cranial elements rather than a true lack of signal; among unmeasured elements within my dataset, high phylogenetic signal in calcaneal gear ratio, associated with foot posture and open versus closed habitat use, has been found for North American carnivorans (Polly et al., 2017), the lengths and shapes of metapodials and phalanges have been found to be highly predictive of arboreality in carnivorans (Van Valkenburgh, 1987), as well as mammals more generally (e.g., Nations et al., 2019), and body shape diversity, as defined by relative measures of the axial skeleton, is consistent with evolution on an adaptive landscape defined by clade specific peaks, albeit peaks that are not obviously associated with ecology (Law, 2021). Further tests of this finding using an expanded ecomorphological dataset are needed to corroborate the lack of early-burst dynamics in locomotor traits for this clade.

While relative sizes and shapes of teeth intuitively relate to food acquisition and processing, the post-cranial skeleton is composed of many moving parts that must work together to facilitate locomotor performance. The post-cranial skeleton therefore tends to exhibit a high degree of inter-element covariance (Young and Hallgŕımsson, 2005; Goswami et al., 2009), which may frustrate the production of discontinuous morphological variation that is maximally optimized for a given locomotor demand (Niklas, 1997, 1999, 2004; Marshall, 2014). Support recovered here for a multi-peak locomotor landscape with relatively low ruggedness, as indicated by a relatively large phylogenetic half-life and equilibrium variances, is consistent with this interpretation and suggests that ecomorphological variation in the carnivoran post-cranial skeleton is far less discontinuous that that found in the mandibulodental system. Scaling of limb proportions in response to changes in body size can also affect functional indices, meaning that more diverse locomotor behaviors can be achieved for a given morphology at small body sizes (Jenkins et al., 1974; Janis and Martín-Serra, 2020; Weaver and Grossnickle, 2020; Wimberly et al., 2021), or that allometric patterns may dominate shape change independent of locomotor behavior at larger sizes (Martín-Serra et al., 2014), further clouding ecomorphological relationships in the post-cranial skeleton. Finally, while locomotor diversity in Carnivora is striking for a mammalian order, it does not span the full breadth of substrate use, locomotor specialization, or morphological diversity encompassed by sequentially higher-level groupings of mammals, such as Laurasiatheria, Boreoeutheria, or Placentalia, where patterns of shape variation associated with substrate use or behavior may be more discontinuous (Chen and Wilson, 2015; Janis and Martín-Serra, 2020; Weaver and Grossnickle, 2020). In other words, failure to detect early-burst like patterns at a given level of the phylogenetic hierarchy, such as in mammalian orders, does not preclude the possibility that a burst occurred at a higher level than the one sampled (Foote, 1996; Jablonski, 2017). The lack of support for partitioning of post-cranial variation among carnivoran clades may well be due to carnivorans themselves sitting atop a rugged peak on a broader mammalian adaptive landscape. A more comprehensive analysis of carnivoran post-cranial morphological evolution is required to refute or confirm the my findings. However, my findings are silent with respect to the hypothesis that mammalian evolution more broadly is characterized by adaptive radiation along an axis of substrate use (Osborn, 1902; Van Valen, 1971), and higher-level tests of this hypothesis are urgently required.

### The Unpredictable Role of Body Size in Macroevolution

Body mass has long been a focal trait for comparative biologists, partly due to its wide availability and ease of access in the literature, but also due to its supposed strong relationship with multiple aspects of life history and ecology (Bonner, 1965; Calder, 1984; Peters, 1986; LaBarbera, 1989). Macroevolutionary studies of diverse clades have yielded support for dynamics that are consistent (Slater et al., 2010; Derryberry et al., 2011; Pyron and Burbrink, 2012) and inconsistent (Harmon et al., 2010; Venditti et al., 2011) with adaptive radiation along an axis of body size variation, but few have explicitly related variation in size to variation in ecological role and, therefore, to macroevolutionary dynamics on the adaptive landscape; indeed, some authors have used the apparently stochastic nature of body mass evolution to argue more generally that trait optima are too ephemeral over macroevolutionary timescales to be a useful paradigm at all in comparative research (Pagel et al., 2022). My results would initially seem to strengthen this conclusion by not only demonstrating that the accumulation of body size disparity within and among carnivoran clades is consistent with a null Brownian motion model (figure 4C; see also Roycroft et al. 2020 for a spatial demonstration of this phenomenon using Australian rodent body mass), but also by showing that there are no pronounced peaks on the body size adaptive landscape. I stress that this result should not be taken to mean that body mass evolution is not adaptive in carnivorans, nor that selection on body mass may not be strong and directional over microevolutionary timescales (Schluter 1988, Kingsolver et al. 2001, Kingsolver and Pfennig 2004, Hereford et al. 2004; but see Rollinson and Rowe 2015). Rather, despite the importance of body mass for many aspects of organismal ecology, microevolutionary patterns need not scale to macroevolutionary levels (Jablonski, 1996), and adaptive peaks associated with body size variation are likely not stable at geographic (Diniz-Filho and T^orres, 2002; Diniz-Filho et al., 2009; Roycroft et al., 2020) or temporal scales ranging from seasonal (Powell and King, 1997) to geologic (Finarelli, 2007; Slater, 2015; Law, 2019).

Still, it is striking that the apparently ephemeral nature of body mass optima contrasts strongly with the stable peaks associated with dietary resource use that have persisted for over 50 million years in this clade (Slater, 2015). This result should not be surprising though, as the material properties of muscle, keratin, and cellulose that influence tooth shape and size are unlikely to have undergone dramatic shifts over this period, even if the climatic and environmental factors or the composition and body size distribution of the prey base that influence carnivoran body mass have systematically varied (Van Valkenburgh, 2007; Slater, 2015).

The decoupling of tempo and mode in mandibulodental and body size evolution recovered here (see also Meloro and Raia, 2010; Slater, 2015; Slater and Friscia, 2019; Grossnickle, 2020) suggests that these traits have played fundamentally different roles in facilitating ecological interactions over macroevolutionary timescales, leading to distinct signals in these comparative data. Instead of providing evidence against the utility of the adaptive landscape metaphor (Pagel et al., 2022), these results together suggest that body size is, in general, likely to be a poor substitute for ecomorphological traits when attempting to understand clade macroevolutionary dynamics (Jablonski, 1996; Slater, 2015; Grossnickle, 2020).

## Conclusion

Adaptive radiation is a provocative term and has been used to describe a range of macroevolutionary scenarios (Givnish, 1997, 2015; Olson and Arroyo-Santos, 2009; Losos et al., 2010), from replicated ecomorph origins in isolated populations of a single species (Schluter, 1993, 1995) to rapid rates of speciation (Slowinski and Guyer, 1989) or phenotypic evolution (Harmon et al., 2010). Despite this diversity of perspectives, using the original formulations of adaptive radiation (Osborn, 1902; Simpson, 1953) we are able to make clear predictions about the structure of ecomorphological variation across phylogeny and the topography of the underlying adaptive landscape on which this variation evolves (Arnold et al., 2001). Evaluating these hypotheses across trait complexes in a single clade can yield novel insights into whether adaptive radiations tend to occur simultaneously across trait complexes (Osborn, 1902), sequentially along distinct axes of resource use (Diamond, 1986; Streelman and Danley, 2003), or, as I have found here, whether some ecological axes are more accessible to diversifying clades than others.

Importantly, while the phylogenetic comparative methods toolkit continues to expand, it is mature enough now that the limitations to addressing questions about the frequency and form of adaptive radiation lie not in the availability of methods but, rather, in the availability of suitable data. My results confirm that body size, while widely accessible and intrinsically interesting in its own right, is an axis of trait variation that evolution is able to exploit rapidly in response to ephemeral shifts in the topography of the adaptive landscape, and there is no guarantee that variation in size is related to niche dynamics that are important over macroevolutionary timescales (Uyeda et al., 2011). I suggest that the most fruitful inference of adaptive radiation dynamics will emerge from analyses of morphological traits that can be directly and explicitly related to the distinct ecological roles that organisms play in their environments. It is time for comparative biologists to move beyond body size and to employ a richer suite of ecomorphological data to more directly evaluate hypotheses about the nature of adaptive radiation.

