## Supplementary File C for "Topographically distinct adaptive landscapes for teeth, skeletons, and size explain the adaptive radiation of Carnivora (Mammalia)"

### Supplementary File C: Analysis of Ecomorphological Diversification

In this document, we will run all the analyses of carnivoran mandibulodental, post-cranial, and body size evolution performed in the manuscript. There are a lot of analyses so we will run them all on each data type before moving on to the next. Comparisons across traits will occur at the end of the document.

We'll begin by loading required packages and custom functions

```
library(geiger)
library(phytools)
library(MASS)
library(mvMORPH)
library(geomorph)
library(motmot)
library(PhylogeneticEM)
library(RColorBrewer)
source("functions_postcranial.R")
source("CustomPhyloEM.R")
```

We can also read in the taxonomic information (family assignment) and the complete carnivoran phylogeny from Slater and Friscia 2019.

```
family <- read.csv("family.csv", stringsAsFactors = F, row.names = 1)
family<-setNames(family[,1], rownames(family))
phy<-ladderize(read.tree("mcc.tre"))
```

This phylogeny can be pruned as needed for each set of analyses.

#### Post-cranial Traits

We begin by reading in the post-cranial traits and pruning the phylogeny to match the data

```
postcranial <- read.csv("meantraitvals.csv", row.names = 1, stringsAsFactors = F)
td<-treedata(phy, postcranial, sort = T, warnings = F)
pc.phy <- td$phy
postcranial <-td$data
rm(td)
```

##### log-shape variables

One point we need to remember is that these traits incorporate size, while we are interested in "shape". We will remove size by computing Mossiman shape variables, or log-shape variables. To do this, we first compute the geometric mean trait value for each taxon, we then divide each trait value by that mean, and then we log transform those values

```
geo.mean<-(apply(postcranial, 1, function(x) prod(x)^(1/length(x)) ))
for(i in 1:nrow(postcranial)) postcranial[i,] <- log(postcranial[i,] / geo.mean[i])
```

#### PCA

Now that we have removed size from the traits, we can examine axes of maximal shape (co)variation using principal components analysis. The scale and variance among traits is quite variable still so we will perform PCA on the correlation, rather than covariance matrix.

```
#ppca.mosiman.pc <-phyl.pca(tree=pc.phy, Y=(postcranial), mode = "corr")
ppca.mosiman.pc <-prcomp(postcranial, scale=T)
var.expl.pc <-round(100*((ppca.mosiman.pc$sdev^2) / sum(ppca.mosiman.pc$sdev^2)),2)
var.expl.pc
```

```
##  [1] 36.88 15.66 10.67  6.63  5.71  4.42  3.28  2.65  2.16  2.08  1.70  1.59
## [13]  1.14  1.05  0.86  0.78  0.69  0.57  0.48  0.37  0.30  0.28  0.06  0.00
```

```
round(ppca.mosiman.pc$sdev^2,2)
```

```
##  [1] 8.85 3.76 2.56 1.59 1.37 1.06 0.79 0.64 0.52 0.50 0.41 0.38 0.27 0.25 0.21
## [16] 0.19 0.17 0.14 0.12 0.09 0.07 0.07 0.01 0.00
```

```
cumsum(var.expl.pc)
```

```
##  [1]  36.88  52.54  63.21  69.84  75.55  79.97  83.25  85.90  88.06  90.14
## [11]  91.84  93.43  94.57  95.62  96.48  97.26  97.95  98.52  99.00  99.37
## [21]  99.67  99.95 100.01 100.01
```

```
broken.stick(var.expl.pc/100)
```

```
## [1] "the first pc to explains less variance than expected is PC4"
```

```
##  [1] 0.157331591 0.115664924 0.094831591 0.080942702 0.070526035 0.062192702
##  [7] 0.055248257 0.049295876 0.044087543 0.039457913 0.035291247 0.031503368
## [13] 0.028031146 0.024826018 0.021849827 0.019072049 0.016467883 0.014016902
## [19] 0.011702087 0.009509105 0.007425772 0.005441645 0.003547705 0.001736111
```

PCs 1-10 collectively account for 90% of the variance and so will be retained for analysis here. Each has an eigenvalue greater than or equal to 0.5, which means that they each account for at least as much variation as a half an individual measurement and each accounts individually for $>$ 2% of the variance

Ultimately we want to plot the PCs. We need to set up colors for each family first:

```
pc.family<-family[rownames(ppca.mosiman.pc$x)]
pc.family<-as.factor(pc.family)
family.col<-c("black",brewer.pal(12, "Paired"))
```

We can test for phylogenetic signal in the PCs using the K mult statistic.

```
physig_postcranial<-physignal(ppca.mosiman.pc$x[,1:3], phy = pc.phy,iter = 9999)
par(mfrow=c(1,1))
hist(physig_postcranial$random.K, breaks=100, col="lightblue",
     main=paste("postcranial Kmult=",
    round(physig_postcranial$phy.signal,2) ), xlab="K")
arrows(x0 = physig_postcranial$phy.signal, y0 = 100,
       x1 = physig_postcranial$phy.signal,
       y1 = 0, col="red", lwd=2)
```

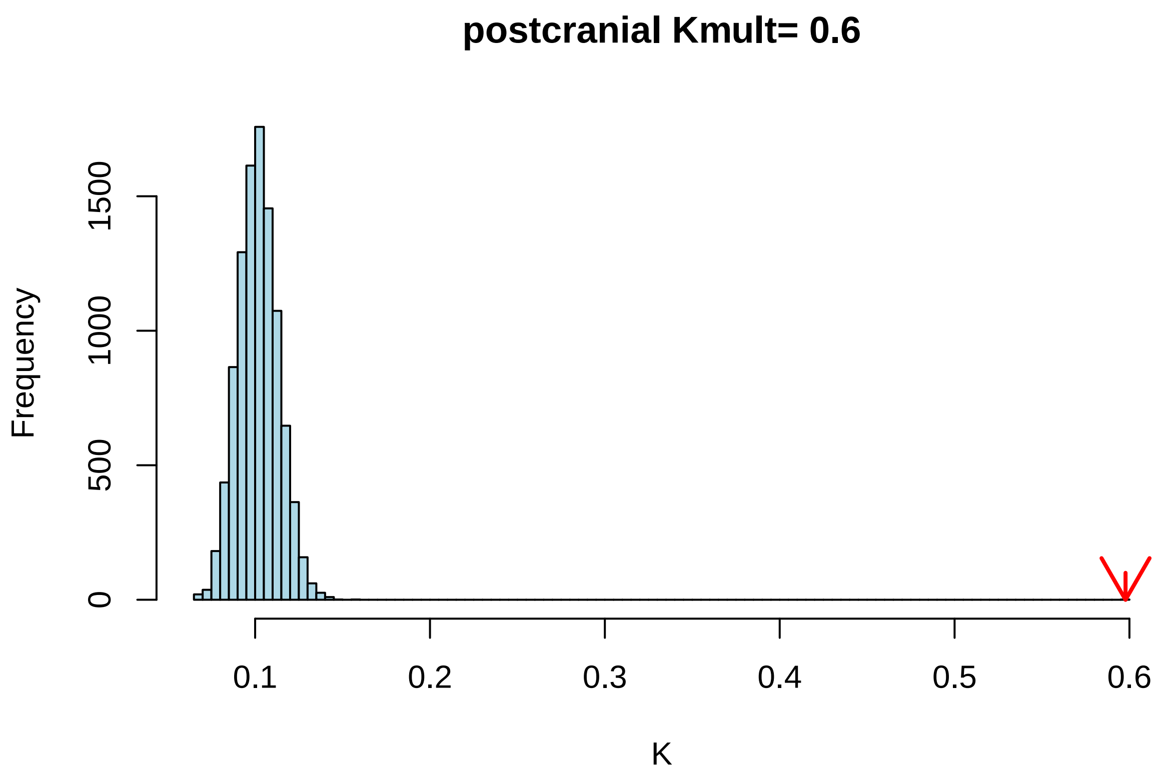


This shows that there is more phylogenetic signal than would be expected from random traits, but K mult ~ 0.5 is still pretty low. We'll interrogate this further.

#### Disparity through time

Disparity through time examines the rate at which trait variation is partitioned among clades, relative to expectation under a constant rates process.

```
pc.dtt<-geiger:::dtt(pc.phy, ppca.mosiman.pc$x[,1:3],nsim = 9999,calculateMDIp = TRUE)
```

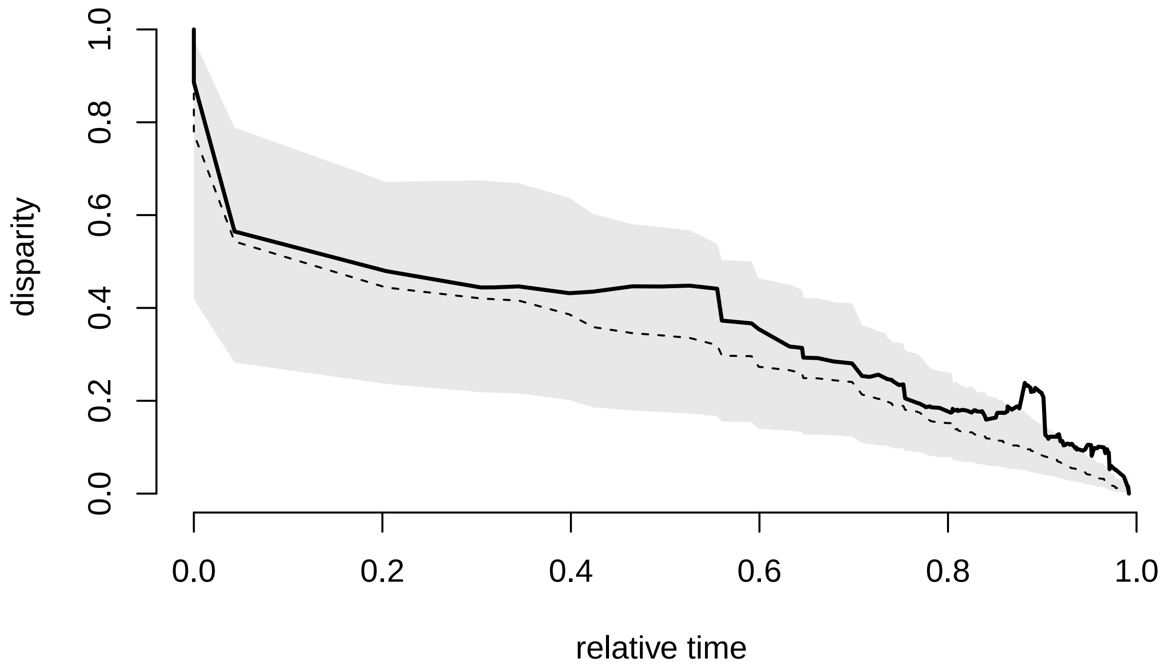


The post-cranial curve is higher than the BM simulations, suggesting a very un-EB like dynamic. We can also use the DTT output to calculate the center of gravity - a weighted average disparity that tells us the point in clade history at which subclade disparity declines to half its total. By comparing this value to the centers of gravity for each BM simulated dataset, we can ask whether trait evolution differs significantly from the NULL expecation. This is something like a multivariate version of the early burst model but, importantly, does not assume process - its just assessing the outcome.

```
pccog<-sum(pc.dtt$dtt*pc.dtt$times) / sum(pc.dtt$dtt)
sim.cog<-apply(pc.dtt$sim, 2, function(x) sum(x*pc.dtt$times) / sum(x))
hist(sim.cog, col="lightblue", main=paste("PCs 1:3 COG =", round(pccog, 2)), xlim=c(0,1))
arrows(x0 = pccog, y0 = 1500,x1 = pccog, y1 = 0,lwd=2, col="red")
```

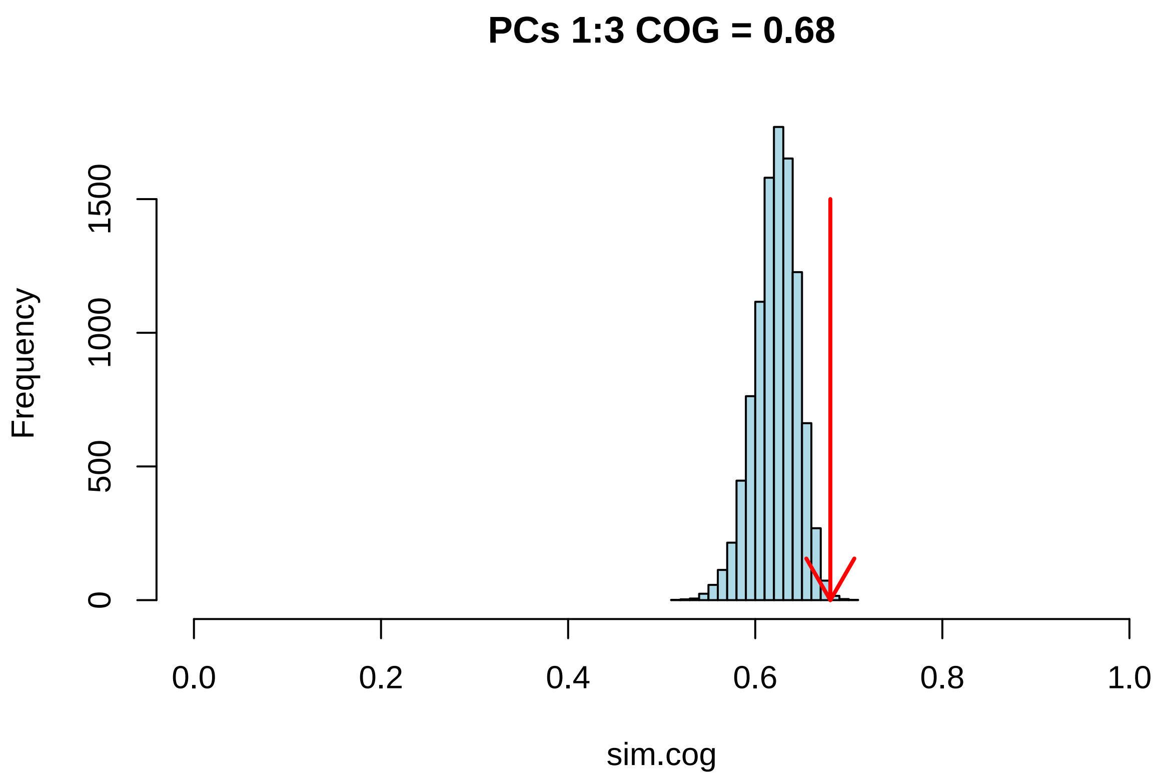


The post-cranial COG (red arrow) is higher (closer to the present) than the BM simuations (blue bars). We can assess significance of this pattern by computing the proportion of simulations with a higher COG than the empirical data:

```
pccog
```

```
## [1] 0.6802723
```

```
length(which(sim.cog>pccog)) / (length(sim.cog))
```

```
## [1] 0.00210021
```

Clearly, post-cranial traits have a significantly higher COG than expected from traits evolving under a constant rates process.

##### Evolution on the post-cranial adaptive landscape

Adaptive radiation occurs when lineages invade distinct adaptive zones, or regions of the adaptive landscape. The signal of these invasions may be present in trait data. We will use a method that is a priori naive to the location and frequency (if at all) of these shifts to identify any that exist. We will allow for negative alphas, consistent with EB dynamics

```
alphas <- find_grid_alpha(phy = pc.phy, nbr_alpha = 100, factor_up_alpha = 1, allow_negative = F)
pc_em_res <- PhyloEM(phylo = pc.phy,
                         Y_data = t(ppca.mosiman.pc$x[,1:3]),
                         process = "scOU",
                         independent = F,
                         random.root = FALSE,                 ## Root is stationary
                         stationary.root = TRUE,
```

```
                         K_max = 30,                          ## Masimal number of shifts
                         parallel_alpha = TRUE,               ## This can be set to TRUE for
                         Ncores = 10,
                         alpha = alphas,
                         allow_negative=FALSE,
                         Nbr_It_Max = 1000)                          ## parallel computations
```

```
## There are some equivalent solutions to the set of shifts selected by the BGHml method.

## There are some equivalent solutions to the set of shifts selected by the BGHlsqraw method.

## There are some equivalent solutions to the set of shifts selected by the BGHmlraw method.
```

```
pc_em_res # no shifts
```

```
## Result of the PhyloEM algorithm.
## Selected parameters by the default method:

## Warning in params_process.PhyloEM(x): There are several equivalent solutions for
## this shift position.

##
## 3 dimensional scOU process with a fixed root.
##
##
## Root value:
## [1]  1.1103016 -0.9508750 -0.7523433
##
## Process variance:
##             [,1]      [,2]       [,3]
## [1,] 0.73738562 0.1976623 0.01732701
## [2,] 0.19766228 0.3921344 0.02440470
## [3,] 0.01732701 0.0244047 0.35190492
##
## Process selection strength:
##           [,1]      [,2]      [,3]
## [1,] 0.1608503 0.0000000 0.0000000
## [2,] 0.0000000 0.1608503 0.0000000
## [3,] 0.0000000 0.0000000 0.1608503
##
## Process root optimal values:
## [1]  1.1103016 -0.9508750 -0.7523433
##
## Shifts positions on branches: 81, 142, 101, 183, 13, 90, 175, 217, 110, 58
## Shifts values:
##            81         142       101       183         13        90         175
## [1,] 3.907510 -1.85599979 -3.368529 -3.383470  4.4087300  3.607338 -1.1228168
## [2,] 1.663046  0.02241331  3.485935  0.232476 -0.0295732  1.122427  5.7342232
## [3,] 1.673267  2.92705568  1.163979  2.480596  3.1442176 -1.239363 -0.3150725
##             217       110        58
## [1,] -5.0674352 -9.227199 5.8871307
## [2,]  0.4364580  2.286666 4.5846174
## [3,]  0.2101014 -4.047296 0.6443438
##
##
##
## See help to see all plotting and handling functions.
```

This model fit finds support for 10 shifts in the OU process mean across Carnivora. The alpha parameter of the OU process results in a phylogenetic half-life of ~ 4 myr, which is consistent with fairly fast rates of adaptation.

```
resParams_pc<-params_process(pc_em_res)
```

```
## Warning in params_process.PhyloEM(pc_em_res): There are several equivalent
## solutions for this shift position.
```

```
 resParams_pc$shifts
```

```
## $edges
##  [1]  81 142 101 183  13  90 175 217 110  58
##
## $values
##          [,1]        [,2]      [,3]      [,4]       [,5]      [,6]       [,7]
## [1,] 3.907510 -1.85599979 -3.368529 -3.383470  4.4087300  3.607338 -1.1228168
## [2,] 1.663046  0.02241331  3.485935  0.232476 -0.0295732  1.122427  5.7342232
## [3,] 1.673267  2.92705568  1.163979  2.480596  3.1442176 -1.239363 -0.3150725
##            [,8]      [,9]     [,10]
## [1,] -5.0674352 -9.227199 5.8871307
## [2,]  0.4364580  2.286666 4.5846174
## [3,]  0.2101014 -4.047296 0.6443438
##
## $relativeTimes
##  [1] 0 0 0 0 0 0 0 0 0 0
```

```
#
CustomPhyloEM(pc.phy, resParams_pc, cex=0.1, edge.width = 2)
```

```
## Loading required package: viridisLite
```

```
##
## Attaching package: 'viridis'
```

```
## The following object is masked from 'package:maps':
##
##     unemp
```

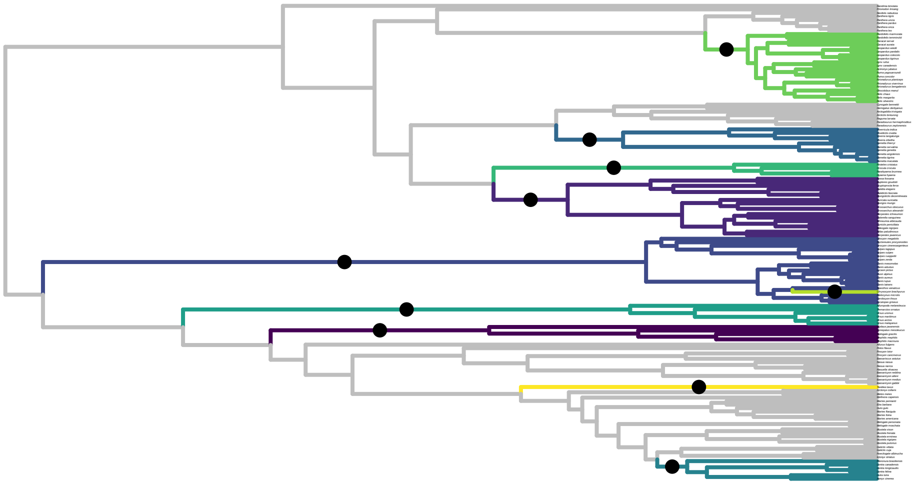

```
eq.shifts.pc<-equivalent_shifts(pc.phy, resParams_pc)
```

we can look at how many equivalent shift configurations there are and calculate support for those in the returned model as the proportion of cases in which that shift occurs

```
shift.freq<- table(eq.shifts.pc$eq_shifts_edges) /ncol(eq.shifts.pc$eq_shifts_edges)
plot(pc.phy, cex=0.3)
edgelabels(edge = as.numeric(names(shift.freq)), text=round(shift.freq,2), cex=0.5)
```

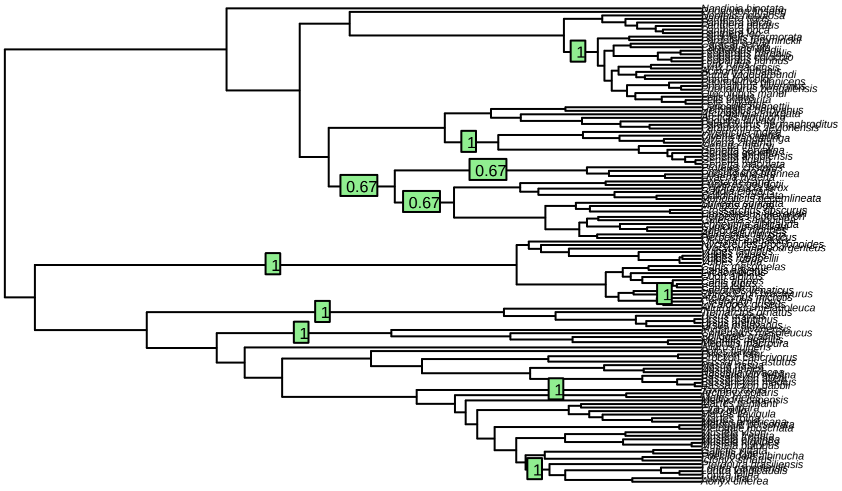

#### Craniodental Evolution

We will now replicate the analyses above for 28 craniodental traits for 198 species from Slater and Friscia (2019). That paper used functional ratios but we will here use the raw linear measurements. As with post-cranial trait evolution, we will remove size by computing log-shape variables based on geometric mean standardization.

```
cranio <- read.csv("craniodental.csv", row.names = 1, stringsAsFactors = F)
# add 1 to all molar traits with 0 for log transformation
cranio[,which(!apply(cranio, 2, function(x)
  sum(which(x==0)))==0)] <- cranio[,which(!apply(cranio, 2, function(x)
      sum(which(x==0)))==0)]+1
geo.mean.cranio<-(apply(cranio, 1, function(x) prod(x)^(1/length(x)) ))
for(i in 1:nrow(cranio)) cranio[i,] <- log(cranio[i,] / geo.mean.cranio[i])

td<-treedata(phy, cranio, sort=T, warnings =F)
cranio <- td$data
cranio.phy<-td$phy
```

##### Principal Components Analysis

We will begin by performing a phylogenetic PCA on the craniodental traits.

```
#ppca.cranio <-phyl.pca(tree=cranio.phy, Y=cranio, mode="cor")
ppca.cranio <- prcomp(x = cranio, scale=TRUE)
var.expl.cranio<- round(100*((ppca.cranio$sdev^2) / sum(ppca.cranio$sdev^2)),2)
```

```
var.expl.cranio
```

```
##  [1] 63.84 12.47  6.76  4.98  2.78  1.94  1.69  1.04  0.88  0.72  0.63  0.45
## [13]  0.45  0.32  0.21  0.16  0.14  0.10  0.10  0.08  0.06  0.05  0.04  0.03
## [25]  0.02  0.02  0.01  0.00
```

```
cumsum(var.expl.cranio)
```

```
##  [1] 63.84 76.31 83.07 88.05 90.83 92.77 94.46 95.50 96.38 97.10 97.73 98.18
## [13] 98.63 98.95 99.16 99.32 99.46 99.56 99.66 99.74 99.80 99.85 99.89 99.92
## [25] 99.94 99.96 99.97 99.97
```

```
round(ppca.cranio$sdev^2,2)
```

```
##  [1] 17.88  3.49  1.89  1.40  0.78  0.54  0.47  0.29  0.25  0.20  0.18  0.13
## [13]  0.13  0.09  0.06  0.05  0.04  0.03  0.03  0.02  0.02  0.01  0.01  0.01
## [25]  0.01  0.01  0.00  0.00
```

```
broken.stick(var.expl.cranio/100)
```

```
## [1] "the first pc to explains less variance than expected is PC3"
```

```
##  [1] 0.140256109 0.104541823 0.086684680 0.074779918 0.065851347 0.058708489
##  [7] 0.052756109 0.047654068 0.043189782 0.039221528 0.035650099 0.032403346
## [13] 0.029427156 0.026679903 0.024128883 0.021747930 0.019515787 0.017414947
## [19] 0.015430820 0.013551121 0.011765406 0.010064726 0.008441350 0.006888555
## [25] 0.005400459 0.003971888 0.002598262 0.001275510
```

5 PCs account for 90% of the variance in dental traits. Each has an eigenvalue >0.5 and indivdiually explains >2% of the variance, making this set comparable in all ways explained to the post-cranial dataset

And again we'll set up some colors for plotting

```
cranio.family<-family[rownames(ppca.cranio$x)]
cranio.family<-as.factor(cranio.family)
```

And now we can plot

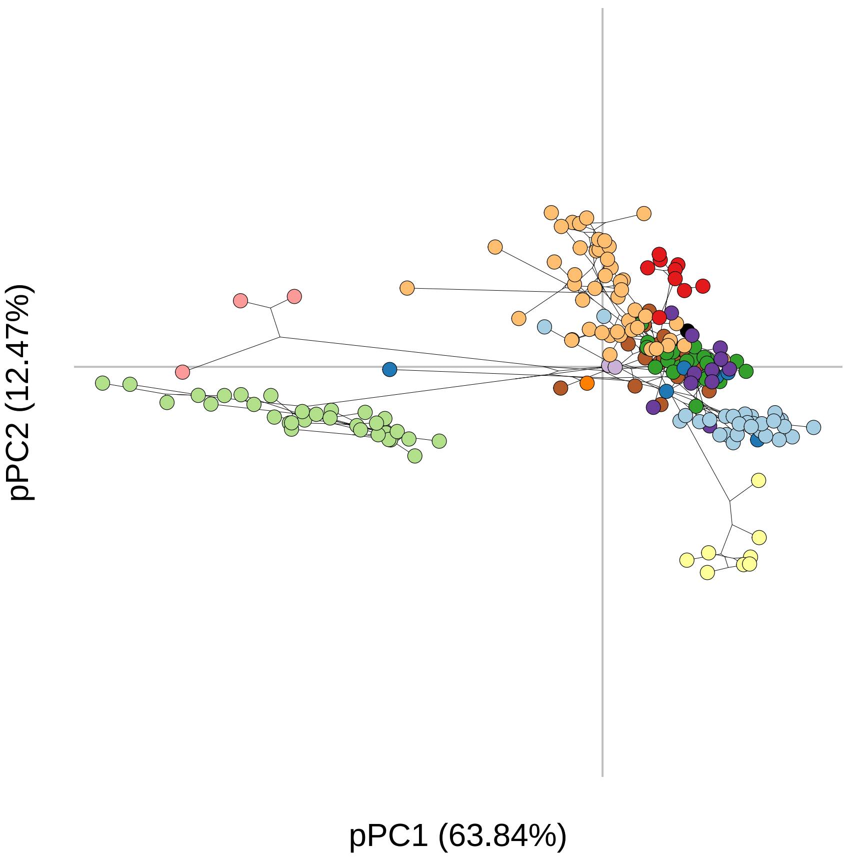


```
## NULL
```

It's clear here that there is very strong phylgenetic signal in these data, with lots of convergence too. Traits associated with carnassial blade length, jaw muscle lever arms and mandible bending strength load negatively on PC1 and correspond to hypercarnivory, while traits associated with molar grinding area load positively - canids occupy an intermediate position on PC1 and ursids etc load positively. PC2 separates taxa with large molars and strong jaws (ursids, procyonids etc) from other taxa.

For the manuscript, we will generate a plot of PC1 vs 2 for both postcranial and craniodental traits, for comparative purposes.

```
pdf(file="Figures/prcomp_comparison.pdf", width=7, height=2)
#par(mfrow=c(1,2),mar=c(3,3,1,1))


nf <- layout(matrix(c(1,2,3), 1, 3, byrow = TRUE),
      widths=c(lcm(3*2.54), lcm(1*2.54),lcm(3*2.54)),
      heights = c(lcm(2*2.54), lcm(2*2.54),lcm(2*2.54)), respect = TRUE)
par(mar=c(3,3,1,1))

customPhylomorph(ppca.cranio$x[,1:2], cranio.phy,
                 xlab=paste("PC1 (", var.expl.cranio[1], "%)", sep=""),
                 ylab=paste("PC2 (", var.expl.cranio[2], "%)", sep=""),
                 pch.cex = 1,color = family.col[cranio.family], phy.lwd = 0.1, axis.cex=0.75, asp=1)
par(xpd=NA)
```

```
mtext(side=2, line=1, text = "A", las=2,at = par("usr")[4]+2, cex=1)
par(xpd=NA)

plot(ppca.mosiman.pc$x[,c(1,2)], type="n", axes=F, xlab="", ylab="")
legend("right", legend=levels(pc.family), pch=21,
       pt.bg =family.col, bty="n", cex=1,pt.cex=1.5)
par(xpd=F)

customPhylomorph(ppca.mosiman.pc$x[,1:2], pc.phy,
                 xlab=paste("PC1 (", var.expl.pc[1], "%)",
                 sep=""), ylab=paste("PC2 (", var.expl.pc[2], "%)", sep=""),
                 pch.cex = 1,color = family.col[pc.family], phy.lwd = 0.1, axis.cex = 0.75, asp=1)
#legend("topleft", legend ="A)", bty="n", cex=2)
mtext(side=2, line=1, text = "B", las=2,at = par("usr")[4], cex=1)
par(xpd=NA)
dev.off()
```

```
## pdf
##   2
```

#### Phylogenetic signal

It looks like there is plenty of phylogenetic signal in the craniodental data, in contrast with the postcranial data. We can test that again using the K mult statistic.

```
physig_craniodental<-physignal(ppca.cranio$x[,1:2], phy = cranio.phy,iter = 9999)
par(mfrow=c(1,1))
hist(physig_craniodental$random.K, breaks=100, col="lightblue",
     main=paste("craniodental Kmult=", round(physig_craniodental$phy.signal,2) ),
     xlab="K")
arrows(x0 = physig_craniodental$phy.signal, y0 = 100,
       x1 = physig_craniodental$phy.signal, y1 = 0,
       col="red", lwd=2)
```

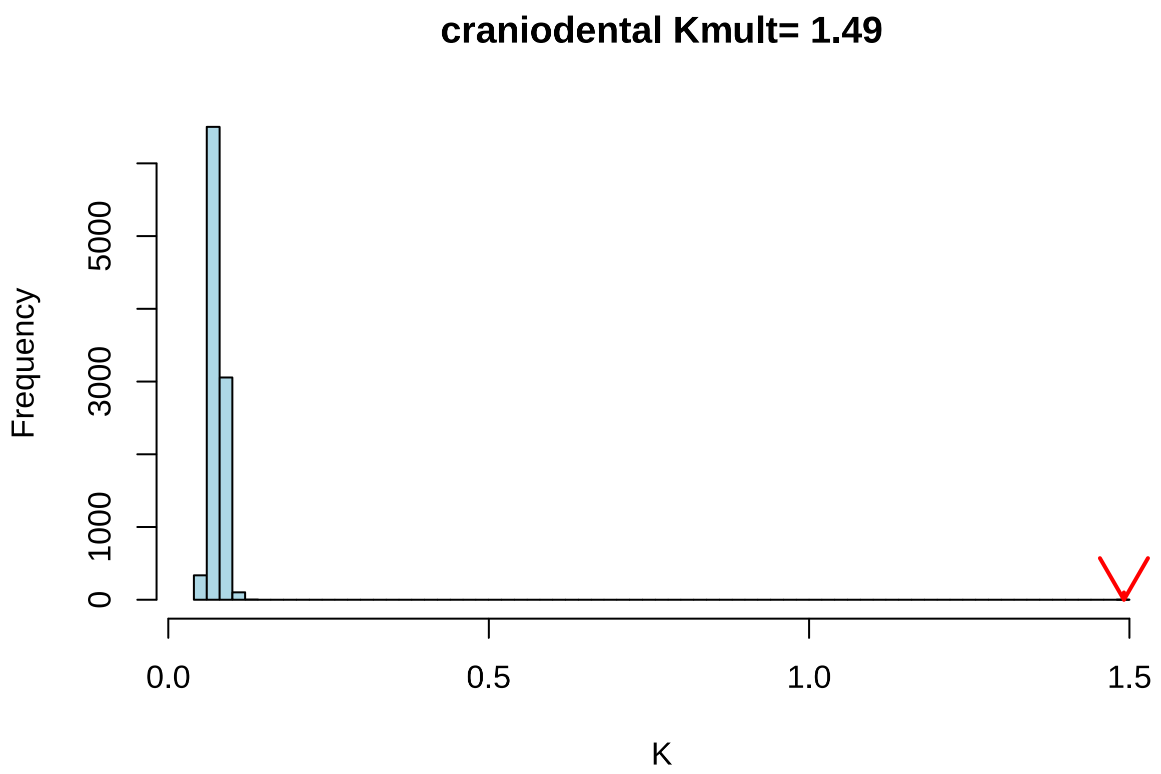


Phylogenetic signal is really high - greater than 1 - for the craniodental traits, which suggests EB-like dynamics are plausible. Slater and Friscia 2019 found early bursts for some traits but not others, but they did not look at multivariate trait evolution. We can test this using the disparity through time approach.

#### Disparity through Time

As before, we can look at the disparity through time curve for the first 4 craniodental PCs to see if parititioning of variation among clades occurs early, midway, or late and compare it to expectations of a constant rates process

```
cranio.dtt<-geiger:::dtt(cranio.phy, ppca.cranio$x[,1:2],nsim = 9999,calculateMDIp = TRUE)
```

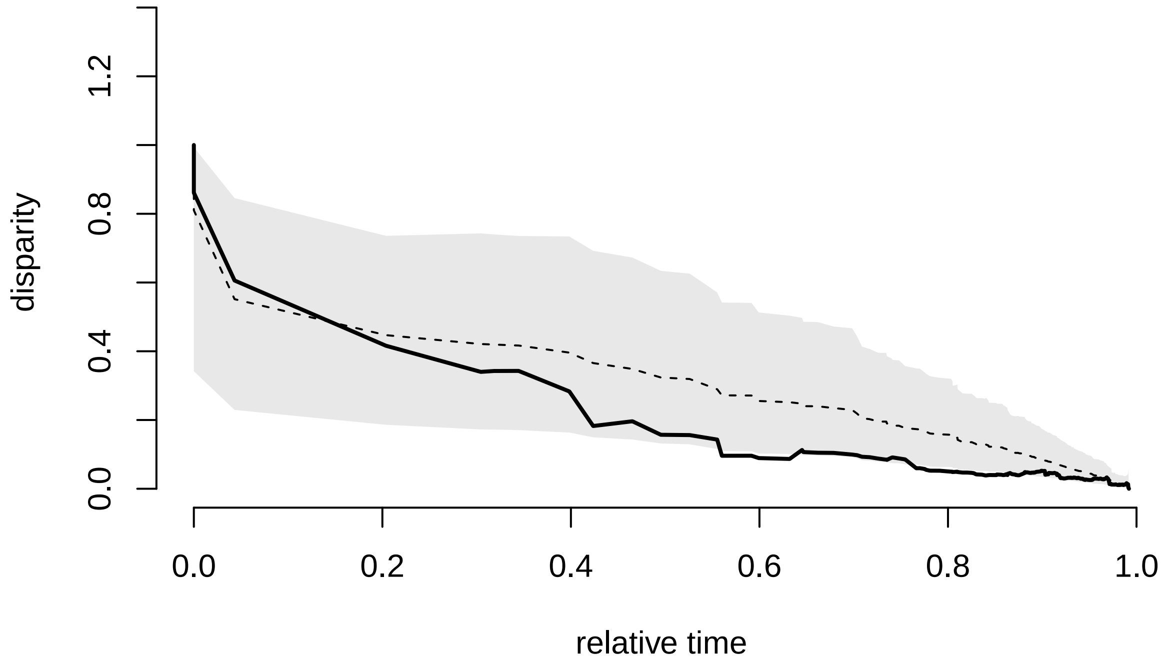


The craniodental curve is mostly lower than the BM simulations, suggesting EB like dynamics. We can again calculate the center of gravity of the curve and compare it to expectations under BM

```
craniocog<-sum(cranio.dtt$dtt*cranio.dtt$times) / sum(cranio.dtt$dtt)
cranio.sim.cog<-apply(cranio.dtt$sim, 2, function(x) sum(x*cranio.dtt$times) / sum(x))
hist(cranio.sim.cog, col="lightblue",
     main=paste("PCs 1:2 COG =", round(craniocog, 2)), xlim=c(0,1))
arrows(x0 = craniocog, y0 = 1500,x1 = craniocog, y1 = 0,lwd=2, col="red")
```

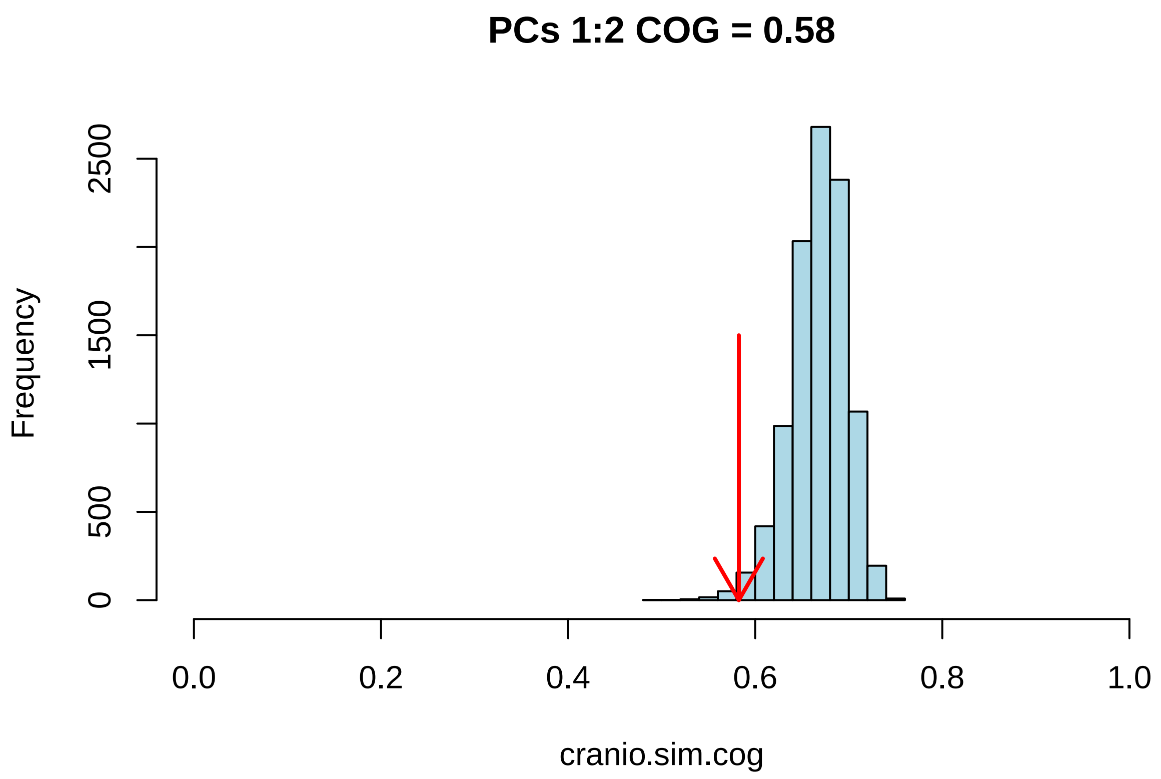


```
quantile(cranio.sim.cog, c(0.025,0.975))
```

```
##      2.5%     97.5%
## 0.6019316 0.7177286
```

So the center of gravity is low, and we can assess significance ...

```
craniocog
```

```
## [1] 0.5824819
```

```
length(which(cranio.sim.cog<craniocog)) / (length(cranio.sim.cog))
```

```
## [1] 0.00840084
```

So craniodental traits have a significantly (p=0.02) lower COG than expected from traits evolving under a constant rates process consistent with EB dynamics, though we need make no assumption about process(es).

#### Craniodental Adaptive Landscape

EB-like dynamics for craniodental traits suggest that there may be distinct adaptive zones that carnivorans radiated into. We can use the phylogenetic EM method to assess this hypothesis, as before, and estimate shift positions.

```
alphas <- find_grid_alpha(phy = cranio.phy, nbr_alpha = 100, factor_up_alpha = 1, allow_negative = F)
cranio_em_res <- PhyloEM(phylo = cranio.phy,
               Y_data = t(ppca.cranio$x[,1:2]),
               process = "scOU",
               independent = F,
```

```r
                random.root = FALSE,                 ## Root is stationary
                stationary.root = TRUE,
                K_max = 30,                          ## Masimal number of shifts
                parallel_alpha = TRUE,               ## This can be set to TRUE for
                Ncores = 10,
                alpha = alphas,
                allow_negative=FALSE,
                Nbr_It_Max = 1000)
```

```
## There are some equivalent solutions to the set of shifts selected by the BGHlsq method.

## There are some equivalent solutions to the set of shifts selected by the BGHml method.

## There are some equivalent solutions to the set of shifts selected by the BGHlsqraw method.
```

```r
cranio_em_res
```

```
## Result of the PhyloEM algorithm.
## Selected parameters by the default method:

## Warning in params_process.PhyloEM(x): There are several equivalent solutions for
## this shift position.

##
## 2 dimensional scOU process with a fixed root.
##
##
## Root value:
## [1] 2.12465502 0.02763739
##
## Process variance:
##             [,1]        [,2]
## [1,]  0.53028010 -0.06854851
## [2,] -0.06854851  0.13771746
##
## Process selection strength:
##           [,1]      [,2]
## [1,] 0.3402486 0.0000000
## [2,] 0.0000000 0.3402486
##
## Process root optimal values:
## [1] 2.12465502 0.02763739
##
## Shifts positions on branches: 325, 123, 6, 140, 155, 326, 13, 141, 279, 45, 380, 347, 300, 275, 177,
## Shifts values:
##               325        123          6        140        155        326          13
## [1,] -1.87203070 -0.2939364 -2.168088  2.087726  1.643024 -6.811154 -0.3824004
## [2,] -0.01210495  2.4137691  1.831981 -3.090068 -1.599378 -1.641007  1.8098465
##             141        279        45         380        347        300         275
## [1,] -0.7881643 -11.176918  1.277815 -4.9722067 -2.379893 -0.9199679 -8.0451014
## [2,] -2.0649247   1.814453 -1.092652  0.9187854  0.512740  1.2243797 -0.1006629
##            177       188          94       283       277         316        90
## [1,] -6.642398 -4.848160 -5.2510321 -2.587557  2.293292 -3.4543074 -3.073103
## [2,]  3.356869  3.913503  0.2663461 -2.240466 -2.230322 -0.6439098  1.475234
##
##
##
```

```
## See help to see all plotting and handling functions.
```

We find a different pattern here than for post-cranial traits. 23 shifts are recovered in the craniodental data, though $\alpha$ is similar, estimated to be ~0.17382, corresponding to a half-life of 3.9 myr

```
resParams<-params_process(cranio_em_res)
```

```
## Warning in params_process.PhyloEM(cranio_em_res): There are several equivalent
## solutions for this shift position.
```

```
 resParams$shifts
```

```
## $edges
##  [1] 325 123   6 140 155 326  13 141 279  45 380 347 300 275 177 188  94 283 277
## [20] 316  90
##
## $values
##              [,1]       [,2]      [,3]      [,4]      [,5]      [,6]       [,7]
## [1,] -1.87203070 -0.2939364 -2.168088  2.087726  1.643024 -6.811154 -0.3824004
## [2,] -0.01210495  2.4137691  1.831981 -3.090068 -1.599378 -1.641007  1.8098465
##             [,8]       [,9]     [,10]      [,11]     [,12]      [,13]      [,14]
## [1,] -0.7881643 -11.176918  1.277815 -4.9722067 -2.379893 -0.9199679 -8.0451014
## [2,] -2.0649247   1.814453 -1.092652  0.9187854  0.512740  1.2243797 -0.1006629
##           [,15]     [,16]      [,17]     [,18]     [,19]      [,20]     [,21]
## [1,] -6.642398 -4.848160 -5.2510321 -2.587557  2.293292 -3.4543074 -3.073103
## [2,]  3.356869  3.913503  0.2663461 -2.240466 -2.230322 -0.6439098  1.475234
##
## $relativeTimes
##  [1] 0 0 0 0 0 0 0 0 0 0 0 0 0 0 0 0 0 0 0 0 0
```

```
#
CustomPhyloEM(cranio.phy, resParams, cex=0.5, edge.width = 2)
```

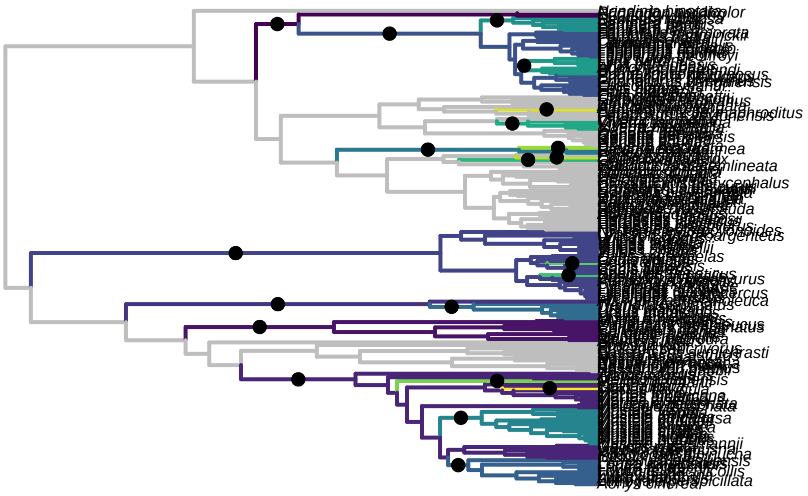

```r
eq.shifts.cranio<-equivalent_shifts(cranio.phy, resParams)
```

we can look at how many equivalent shift configurations there are and calculate support for those in the returned model as the proportion of cases in which that shift occurs

```r
eq.shifts.cranio
```

```
##
## There are 72 equivalent solutions.
##
## Use function plot to see them all.
```

```r
shift.freq<- table(eq.shifts.cranio$eq_shifts_edges) /ncol(eq.shifts.cranio$eq_shifts_edges)
plot(cranio.phy, cex=0.3)
edgelabels(edge = as.numeric(names(shift.freq)), text=round(shift.freq,2), cex=0.5)
```

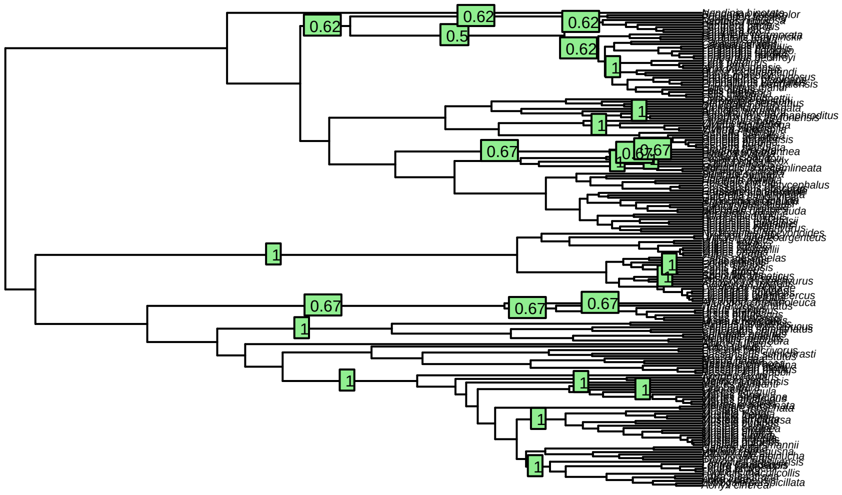

There are over 6000 equivalent shift configurations but they mostly involve the shifting of a more proscribed set of individual shifts (e.g. shifts in one of a pair of sister taxa vs the other).

#### Body mass

We will now do the same set of analyses for body mass. I'll skip through the text a bit because we're repeating ourselves

```
mass <- read.csv("mass.csv", row.names = 1, stringsAsFactors = F)
mass <- setNames(mass[,1], rownames(mass))
td<-treedata(phy, mass, sort = T, warnings = F)
mass.phy <- td$phy
mass <-td$data
rm(td)
```

```
physig_mass<-physignal(mass, mass.phy,iter = 9999)
par(mfrow=c(1,1))
hist(physig_mass$random.K, breaks=100, col="lightblue",
     main=paste("mass Kmult=", round(physig_mass$phy.signal,2) ),
     xlab="K")
arrows(x0 = physig_mass$phy.signal, y0 = 100,
       x1 = physig_mass$phy.signal, y1 = 0, col="red", lwd=2)
```

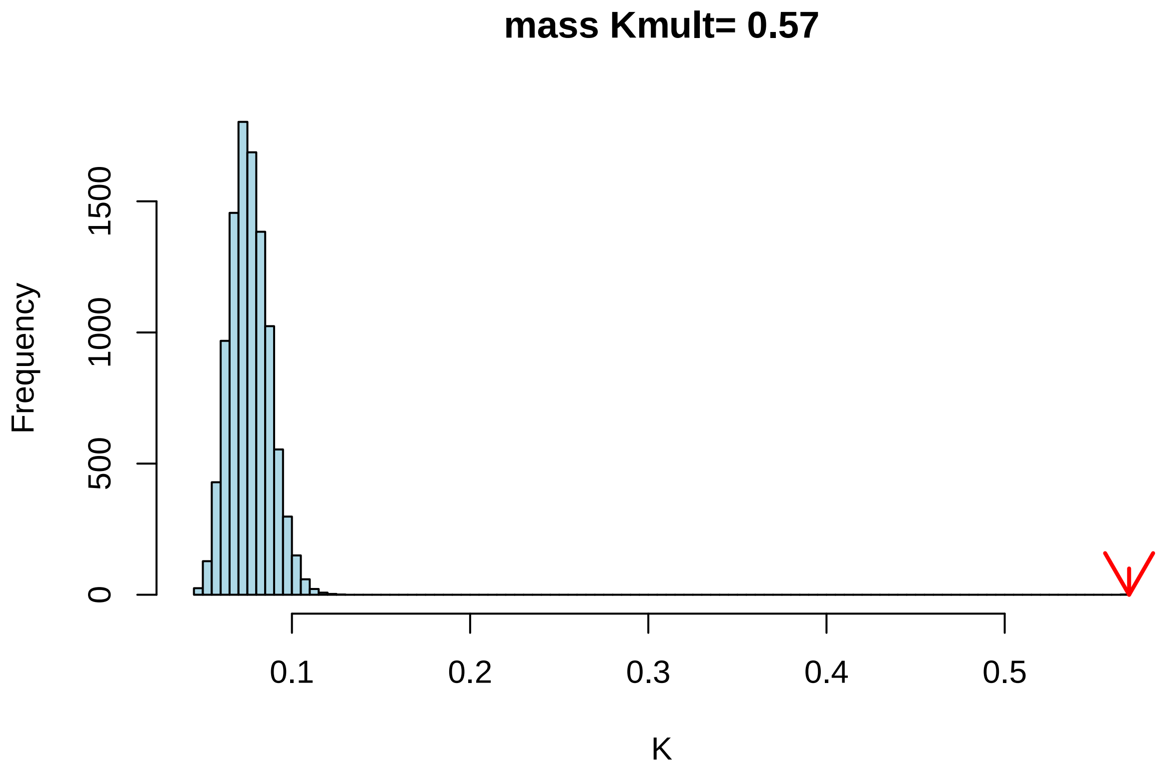


This shows that there is more phylogenetic signal than would be expected from random traits and for post-cranial traits, but K mult lower than for craniodental traits.

##### Disparity through time

```
mass.dtt<-geiger:::dtt(mass.phy, mass,nsim = 9999,calculateMDIp = TRUE)
```

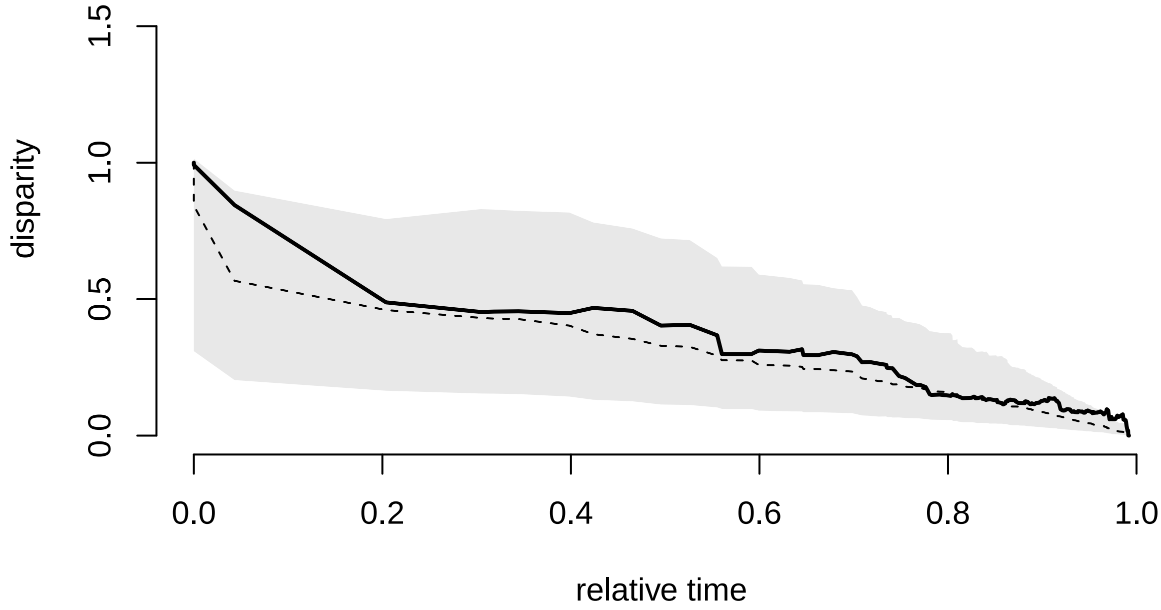


```
masscog<-sum(mass.dtt$dtt*mass.dtt$times) / sum(mass.dtt$dtt)
sim.cog<-apply(mass.dtt$sim, 2, function(x) sum(x*mass.dtt$times) / sum(x))
hist(sim.cog, col="lightblue", main=paste("mass COG =", round(masscog, 2)), xlim=c(0,1))
arrows(x0 = masscog, y0 = 1500,x1 = masscog, y1 = 0,lwd=2, col="red")
```

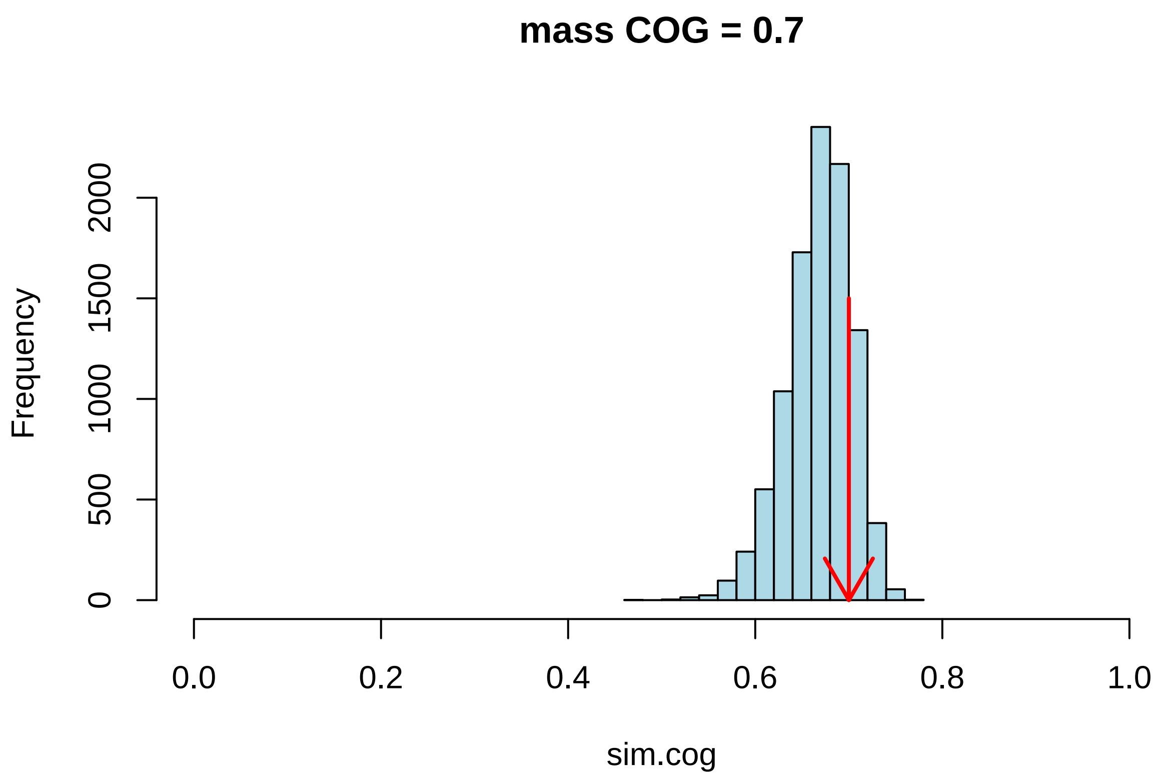


```
masscog
```

```
## [1] 0.7001029
```

```
length(which(sim.cog>masscog)) / (length(sim.cog))
```

```
## [1] 0.1771177
```

the COG for mass is between those of craniodental traits (earlier) and post-cranial traits (later)

##### Evolution on the body mass adaptive landscape

```
alphas <- find_grid_alpha(phy = mass.phy, nbr_alpha = 100, factor_up_alpha = 1, allow_negative = F)
mass_em_res <- PhyloEM(phylo = mass.phy,
               Y_data = t(mass),
               process = "OU", ## not scalar OU model (no covars)
                alpha = alphas,
               random.root = FALSE,                ## Root is stationary
               stationary.root = TRUE,
               K_max = 30,                        ## Masimal number of shifts
               parallel_alpha = TRUE,             ## This can be set to TRUE for
               Ncores = 10,
               Nbr_It_Max = 1000,
               allow_negative = F)                        ## parallel computations
mass_em_res # no shifts
```

```
## Result of the PhyloEM algorithm.
## Selected parameters by the default method:
```

```
## 1 dimensional OU process with a fixed root.
##
##
## Root value:
## [1] 0.5394298
##
## Process variance:
##            [,1]
## [1,] 0.01321722
##
## Process selection strength:
##           [,1]
## [1,] 0.0177146
##
## Process root optimal values:
## [1] 0.5394298
##
## This process has no shift.
##
##
## See help to see all plotting and handling functions.
```

This model fit finds no support for shifts in the OU process mean across Carnivora and alpha is an order of magnitude smaller.

```
## pdf
##   2
```

Table C1: PCA loadings for the first 2 PCs from analysis of craniodental traits.

| | PC1 | PC2 |
|---|---|---|
| p4MD | -0.204 | 0.024 |
| p4BL | -0.198 | 0.159 |
| m1MD | -0.166 | 0.158 |
| m13L | -0.194 | 0.025 |
| m1tL | 0.184 | 0.280 |
| m13BL | -0.100 | 0.200 |
| m1tBL | 0.188 | 0.266 |
| m2MD | 0.225 | 0.076 |
| m2tBL | 0.216 | 0.155 |
| m3MD | 0.115 | -0.357 |
| m3BL | 0.115 | -0.348 |
| MandL | -0.176 | -0.199 |
| MATemp | -0.183 | 0.143 |
| MAMass | -0.166 | -0.016 |
| MDm1m2 | -0.220 | -0.070 |
| MWm1m2 | -0.223 | 0.021 |
| MDp4m1 | -0.210 | -0.150 |
| MWp4m1 | -0.230 | 0.007 |
| MDp3p4 | -0.209 | -0.110 |
| MWp3p4 | -0.226 | 0.032 |
| C1MD | -0.210 | -0.083 |
| CBL | -0.216 | -0.015 |
| P4MD | -0.198 | 0.066 |
| P4BLmx | -0.179 | 0.255 |
| M1MD | 0.170 | 0.185 |
| M1BL | 0.131 | 0.358 |
| M2MD | 0.174 | -0.293 |
| M2BL | 0.174 | -0.234 |

Table C2: PCA loadings for the first 3 PCs from analysis of postcranial traits.

| | PC1 | PC2 | PC3 |
|---|---|---|---|
| scapH | -0.188 | 0.228 | 0.201 |
| scapW | 0.013 | -0.277 | -0.073 |
| glenL | 0.228 | 0.213 | -0.030 |
| glenW | 0.161 | 0.233 | -0.168 |
| HumL | -0.262 | -0.087 | -0.216 |
| HD | 0.206 | -0.019 | -0.272 |
| HEB | 0.297 | -0.074 | -0.125 |
| RL | -0.293 | 0.168 | -0.105 |
| RCL | 0.255 | 0.088 | -0.303 |
| RCB | 0.215 | 0.249 | -0.202 |
| UL | -0.288 | 0.158 | -0.075 |
| UOL | 0.134 | 0.108 | 0.245 |
| FL | -0.281 | -0.105 | -0.191 |
| FW | 0.093 | -0.367 | 0.123 |
| FD | -0.060 | -0.398 | 0.113 |
| FCD | -0.122 | 0.155 | 0.360 |
| FCB | 0.183 | -0.286 | 0.126 |
| FPB | 0.215 | -0.242 | -0.163 |
| MAGT | 0.174 | 0.080 | 0.188 |
| TL | -0.287 | -0.117 | 0.029 |
| PL | 0.095 | -0.218 | 0.393 |
| PSL | -0.196 | -0.087 | -0.136 |
| ischB | 0.058 | 0.276 | 0.322 |
| ilB | 0.200 | 0.059 | 0.172 |
