## Supplementary File A for "Topographically distinct adaptive landscapes for teeth, skeletons, and size explain the adaptive radiation of Carnivora (Mammalia)"

### Supplementary File A: Testing for Locomotor Signal in postcranial indices

#### Linear Discriminant Function Analysis

To investigate whether the functional ratios contain sufficient information to distinguish among carnivorans with distinct locomotor modes and patterns of substrate use, I performed a linear discriminant function analysis using the `lda` function from the `MASS` library. To perform this analysis, we will need the script containing utility functions, as well as csv files containing the trait data and locomotor information:

```
library(MASS)
source("functions_postcranial.R")

## read in the data ##
mean.rat <- read.csv("postcranial_ratios/mean.ratios.csv", row.names=1)
habit <- read.csv("locomotor_mode.csv", stringsAsFactors = F, row.names = 1, header=F)
loc <- setNames(habit[,1], rownames(habit))

## perform dfa ##
dfa<-lda(loc ~.,as.data.frame(mean.rat))
```

We can examine the relative amount of variance explained by each DF axis by calling

```
(dfa$svd^2)/sum(dfa$svd^2)
```

```
## [1] 0.54683244 0.29996473 0.08149963 0.07170320
```

which shows that the first DF axis accounts for around 55 % of the variance while the second is around 30%. To obtain species scores along each function we must multiply each species' trait values by the corresponding scaling factors (Table A1) and then sum them:

```
df1<-apply(mean.rat, 1, function(x) sum(x*dfa$scaling[,1]))
df2<-apply(mean.rat, 1, function(x) sum(x*dfa$scaling[,2]))
plot(df1, df2, pch=21, bg=c("seagreen", "cornflowerblue", "plum3", "salmon", "orange")
     [as.factor(loc)], cex=1.5)
legend("topright", legend=levels(as.factor(loc)), pch=21,
       pt.bg=c("seagreen", "cornflowerblue", "plum3", "salmon", "orange"))
```

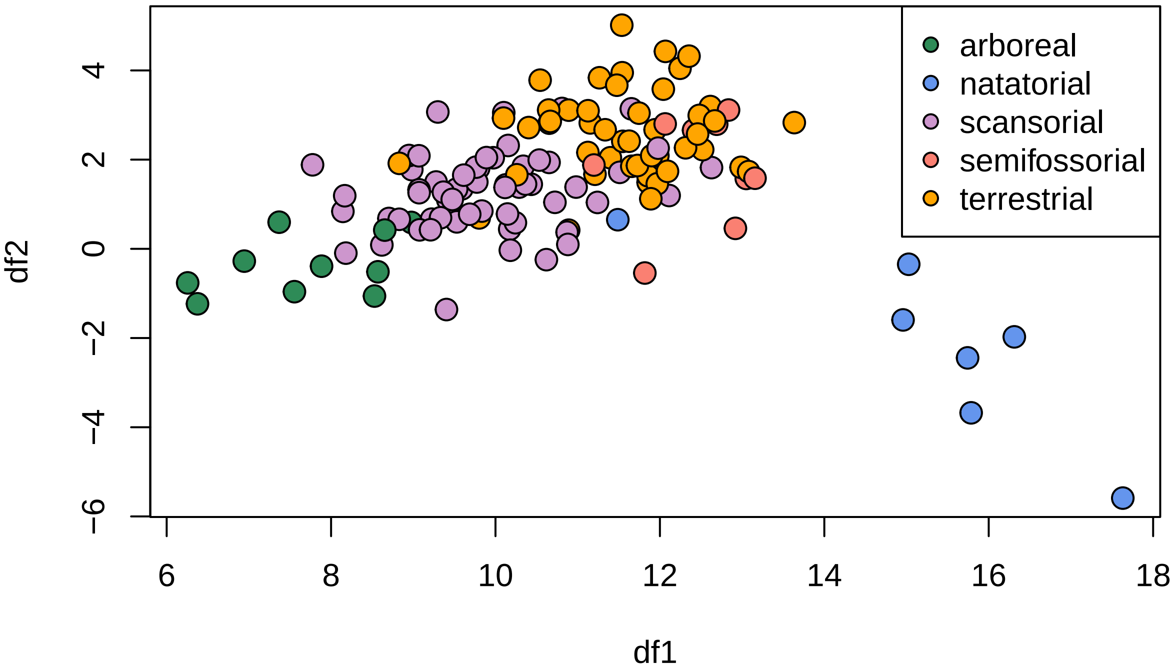


We can now predict group membership for each taxon based on its traits using the discriminant functions. We can also repeat the linear discriminant analysis but with the option CV=TRUE to perform a leave–one–out cross–validation. The aim here is to determine, for each species, whether its combination of trait values yields a reasonable prediction of its group membership given the traits present in other taxa.

```
pred <-MASS:::predict.lda(dfa, as.data.frame(mean.rat))$class
dfa2<-MASS:::lda(loc ~.,as.data.frame(mean.rat), CV=TRUE)
head(cbind(loc, as.character(pred), round(dfa2$posterior,2)))
```

```
##                     loc                         arboreal natatorial scansorial
## Acinonyx_jubatus   "scansorial" "terrestrial" "0"      "0"        "0.24"
## Puma_concolor      "scansorial" "scansorial"  "0.03"   "0"        "0.97"
## Puma_yagouaroundi "scansorial" "scansorial"  "0"      "0"        "0.97"
## Lynx_canadensis    "scansorial" "scansorial"  "0"      "0"        "1"
## Lynx_rufus         "scansorial" "scansorial"  "0"      "0"        "0.98"
## Felis_silvestris   "scansorial" "scansorial"  "0"      "0"        "0.89"
##                     semifossorial terrestrial
## Acinonyx_jubatus   "0"           "0.76"
## Puma_concolor      "0"           "0"
## Puma_yagouaroundi "0"           "0.02"
## Lynx_canadensis    "0"           "0"
## Lynx_rufus         "0"           "0.02"
## Felis_silvestris   "0"           "0.11"
```

In the first 6 rows (which are all felids) we see that the posterior probability that theses species belong to the scansorial category is generally high, though the cheetah is perhaps more similar to terrestrial taxa. The full set of cross–validated assignments are provided in Table A2.

Table A1: Scaling matrix for the first two linear discriminant functions. The magnitude of the scaling factors indicates the importance of each variable on that function while the sign indicates the direction of the effect

| | LD1 | LD2 |
|---|---|---|
| scapula.index | -7.73 | -3.77 |
| glenoid.shape | -7.93 | -0.63 |
| brachial.index | 0.38 | 8.55 |
| humeral.epicondylar.breadth | 4.10 | 2.78 |
| capitulum.shape | 3.18 | 3.09 |
| fossoriality.index | -5.90 | 14.26 |
| crural.index | 2.72 | -9.62 |
| femoral.shaft.shape | 0.76 | 0.21 |
| femoral.epicondylar.width | 48.91 | -26.45 |
| patella.grove.index | -10.27 | 6.18 |
| femoral.epicondylar.index | 13.13 | 208.71 |
| gluteal.index | 17.47 | 1.37 |
| intermembranal.index | 5.54 | -4.80 |
| ischial.breadth | -0.75 | 6.73 |
| iliac.breadth | -0.54 | -1.24 |
| pubic.symphysis.length | 6.95 | 2.89 |

Table A2: Leave–one–out cross–validation of carnivoran locomotor classification. loc gives the assigned locomotor mode, predicted gives the best estimate as to group membership and subsequent columns give the posterior probability that the row taxon belongs to that group.

| | locomotor | predicted | arboreal | natatorial | scansorial | semifossorial | terrestrial |
|---|---|---|---|---|---|---|---|
| Acinonyx jubatus | scansorial | terrestrial | 0 | 0 | 0.24 | 0 | 0.76 |
| Puma concolor | scansorial | scansorial | 0.03 | 0 | 0.97 | 0 | 0 |
| Puma yagouaroundi | scansorial | scansorial | 0 | 0 | 0.97 | 0 | 0.02 |
| Lynx canadensis | scansorial | scansorial | 0 | 0 | 1 | 0 | 0 |
| Lynx rufus | scansorial | scansorial | 0 | 0 | 0.98 | 0 | 0.02 |
| Felis silvestris | scansorial | scansorial | 0 | 0 | 0.89 | 0 | 0.11 |
| Felis margarita | scansorial | scansorial | 0 | 0 | 0.79 | 0 | 0.21 |
| Felis chaus | scansorial | scansorial | 0 | 0 | 0.97 | 0 | 0.03 |
| Otocolobus manul | scansorial | scansorial | 0 | 0 | 0.68 | 0 | 0.31 |
| Prionailurus bengalensis | scansorial | scansorial | 0 | 0 | 0.93 | 0 | 0.07 |
| Prionailurus viverrinus | scansorial | scansorial | 0 | 0 | 0.98 | 0 | 0.01 |
| Prionailurus planiceps | scansorial | scansorial | 0.01 | 0 | 0.97 | 0 | 0.02 |
| Caracal aurata | scansorial | scansorial | 0.02 | 0 | 0.97 | 0 | 0 |
| Caracal serval | scansorial | scansorial | 0 | 0 | 0.99 | 0 | 0.01 |
| Leopardus colocolo | scansorial | scansorial | 0 | 0 | 0.97 | 0 | 0.03 |
| Leopardus tigrinus | scansorial | scansorial | 0 | 0 | 0.95 | 0 | 0.05 |
| Leopardus pardalis | scansorial | scansorial | 0 | 0 | 0.98 | 0 | 0.01 |
| Leopardus wiedii | scansorial | scansorial | 0 | 0 | 0.98 | 0 | 0.02 |
| Pardofelis temminckii | scansorial | scansorial | 0 | 0 | 0.86 | 0 | 0.14 |
| Pardofelis marmorata | scansorial | scansorial | 0.17 | 0 | 0.83 | 0 | 0 |
| Neofelis nebulosa | scansorial | scansorial | 0 | 0 | 0.99 | 0 | 0 |
| Panthera leo | terrestrial | scansorial | 0 | 0 | 0.7 | 0.14 | 0.15 |
| Panthera onca | scansorial | scansorial | 0 | 0 | 0.88 | 0.07 | 0.04 |
| Panthera pardus | scansorial | scansorial | 0 | 0 | 0.98 | 0 | 0.02 |
| Panthera uncia | scansorial | scansorial | 0 | 0 | 0.85 | 0 | 0.15 |
| Panthera tigris | terrestrial | scansorial | 0 | 0 | 1 | 0 | 0 |
| Prionodon linsang | scansorial | scansorial | 0 | 0 | 0.85 | 0 | 0.15 |
| Arctictis binturong | arboreal | arboreal | 0.95 | 0 | 0.05 | 0 | 0 |
| Paguma larvata | scansorial | scansorial | 0.09 | 0 | 0.9 | 0 | 0.01 |
| Paradoxurus hermaphroditus | arboreal | scansorial | 0.29 | 0 | 0.7 | 0 | 0.01 |
| Paradoxurus zeylonensis | arboreal | arboreal | 0.93 | 0 | 0.07 | 0 | 0 |
| Arctogalidia trivirgata | arboreal | arboreal | 0.95 | 0 | 0.05 | 0 | 0 |

| | | | | | | | |
|---|---|---|---|---|---|---|---|
| Hemigalus derbyanus | scansorial | scansorial | 0.04 | 0 | 0.96 | 0 | 0 |
| Cynogale bennettii | scansorial | scansorial | 0 | 0 | 0.92 | 0.01 | 0.07 |
| Civettictis civetta | scansorial | terrestrial | 0 | 0 | 0.02 | 0 | 0.98 |
| Viverra zibetha | scansorial | scansorial | 0 | 0 | 0.81 | 0 | 0.19 |
| Viverra tangalunga | scansorial | scansorial | 0 | 0 | 0.84 | 0 | 0.16 |
| Viverricula indica | scansorial | scansorial | 0 | 0 | 0.74 | 0 | 0.26 |
| Genetta angolensis | scansorial | scansorial | 0.01 | 0 | 0.76 | 0 | 0.24 |
| Genetta maculata | scansorial | scansorial | 0 | 0 | 0.9 | 0 | 0.1 |
| Genetta tigrina | scansorial | scansorial | 0 | 0 | 0.84 | 0 | 0.15 |
| Genetta genetta | scansorial | scansorial | 0 | 0 | 0.73 | 0 | 0.27 |
| Genetta servalina | scansorial | scansorial | 0 | 0 | 0.96 | 0 | 0.04 |
| Genetta thierryi | scansorial | scansorial | 0 | 0 | 0.92 | 0 | 0.08 |
| Atilax paludinosus | natatorial | scansorial | 0 | 0 | 0.68 | 0.01 | 0.31 |
| Herpestes javanicus | terrestrial | terrestrial | 0 | 0 | 0.01 | 0.46 | 0.53 |
| Bdeogale nigripes | terrestrial | terrestrial | 0 | 0 | 0.16 | 0.51 | 0.33 |
| Cynictis penicillata | terrestrial | terrestrial | 0 | 0 | 0 | 0 | 1 |
| Ichneumia albicauda | terrestrial | terrestrial | 0 | 0 | 0.56 | 0 | 0.44 |
| Galerella sanguinea | scansorial | terrestrial | 0 | 0 | 0.04 | 0 | 0.96 |
| Herpestes ichneumon | terrestrial | terrestrial | 0 | 0 | 0.1 | 0.56 | 0.35 |
| Crossarchus alexandri | scansorial | terrestrial | 0 | 0 | 0.13 | 0.15 | 0.73 |
| Crossarchus obscurus | scansorial | terrestrial | 0 | 0 | 0.13 | 0.01 | 0.86 |
| Mungos mungo | terrestrial | terrestrial | 0 | 0 | 0.13 | 0.03 | 0.84 |
| Suricata suricatta | terrestrial | terrestrial | 0 | 0 | 0 | 0 | 1 |
| Cryptoprocta ferox | scansorial | scansorial | 0.14 | 0 | 0.86 | 0 | 0 |
| Galidia elegans | terrestrial | terrestrial | 0 | 0 | 0.28 | 0 | 0.71 |
| Galidictis fasciata | terrestrial | scansorial | 0 | 0 | 0.88 | 0 | 0.12 |
| Mungotictis decemlineata | terrestrial | terrestrial | 0 | 0 | 0.38 | 0 | 0.61 |
| Eupleres goudotii | terrestrial | terrestrial | 0 | 0 | 0.01 | 0.04 | 0.95 |
| Fossa fossana | terrestrial | terrestrial | 0 | 0 | 0.43 | 0 | 0.57 |
| Crocuta crocuta | terrestrial | terrestrial | 0 | 0 | 0.04 | 0.02 | 0.94 |
| Hyaena hyaena | terrestrial | terrestrial | 0 | 0 | 0 | 0.01 | 0.98 |
| Parahyaena brunnea | terrestrial | terrestrial | 0 | 0 | 0.01 | 0.01 | 0.98 |
| Proteles cristatus | terrestrial | terrestrial | 0 | 0 | 0.1 | 0 | 0.9 |
| Nandinia binotata | scansorial | scansorial | 0.06 | 0 | 0.94 | 0 | 0 |
| Ailuropoda melanoleuca | scansorial | arboreal | 1 | 0 | 0 | 0 | 0 |
| Tremarctos ornatus | scansorial | scansorial | 0 | 0 | 0.92 | 0.07 | 0.01 |
| Ursus malayanus | scansorial | scansorial | 0.3 | 0 | 0.54 | 0.11 | 0.05 |
| Ursus arctos | scansorial | scansorial | 0 | 0 | 0.99 | 0 | 0 |
| Ursus maritimus | terrestrial | scansorial | 0.01 | 0 | 0.99 | 0 | 0 |
| Ursus ursinus | scansorial | scansorial | 0 | 0 | 0.81 | 0 | 0.18 |
| Ailurus fulgens | arboreal | arboreal | 0.36 | 0 | 0.63 | 0 | 0.01 |
| Aonyx cinerea | natatorial | natatorial | 0 | 0.93 | 0 | 0.03 | 0.04 |
| Lutra lutra | natatorial | natatorial | 0 | 1 | 0 | 0 | 0 |
| Lontra canadensis | natatorial | natatorial | 0 | 1 | 0 | 0 | 0 |
| Lontra felina | natatorial | natatorial | 0 | 1 | 0 | 0 | 0 |
| Lontra longicaudis | natatorial | natatorial | 0 | 1 | 0 | 0 | 0 |
| Pteronura brasiliensis | natatorial | natatorial | 0 | 1 | 0 | 0 | 0 |
| Galictis cuja | semifossorial | semifossorial | 0 | 0 | 0.14 | 0.07 | 0.78 |
| Galictis vittata | semifossorial | semifossorial | 0 | 0 | 0.01 | 0.93 | 0.06 |
| Ictonyx striatus | scansorial | terrestrial | 0 | 0 | 0.01 | 0.39 | 0.61 |
| Poecilogale albinucha | terrestrial | terrestrial | 0 | 0 | 0.02 | 0.18 | 0.81 |
| Mustela frenata | terrestrial | terrestrial | 0 | 0 | 0.38 | 0.01 | 0.62 |
| Mustela vison | terrestrial | scansorial | 0 | 0 | 0.98 | 0 | 0.02 |
| Mustela putorius | terrestrial | terrestrial | 0 | 0 | 0.54 | 0.03 | 0.43 |
| Mustela nigripes | terrestrial | terrestrial | 0 | 0 | 0.06 | 0.31 | 0.64 |
| Mustela erminea | terrestrial | terrestrial | 0 | 0 | 0.1 | 0 | 0.89 |
| Melogale moschata | semifossorial | semifossorial | 0 | 0 | 0.56 | 0.15 | 0.29 |
| Melogale personata | semifossorial | semifossorial | 0 | 0 | 0.01 | 0.89 | 0.1 |
| Eira barbara | scansorial | scansorial | 0 | 0 | 0.97 | 0.02 | 0.02 |
| Martes pennanti | scansorial | scansorial | 0 | 0 | 0.95 | 0.02 | 0.04 |
| Gulo gulo | semifossorial | scansorial | 0 | 0 | 0.88 | 0.01 | 0.1 |
| Martes americana | scansorial | scansorial | 0 | 0 | 0.95 | 0 | 0.04 |
| Martes foina | scansorial | scansorial | 0 | 0 | 0.97 | 0.01 | 0.02 |
| Martes flavigula | scansorial | scansorial | 0.12 | 0 | 0.87 | 0 | 0 |

| | | | | | | | |
|---|---|---|---|---|---|---|---|
| Mellivora capensis | semifossorial | semifossorial | 0 | 0 | 0.05 | 0.72 | 0.24 |
| Arctonyx collaris | semifossorial | semifossorial | 0 | 0 | 0.01 | 0.66 | 0.33 |
| Meles meles | semifossorial | semifossorial | 0 | 0 | 0.04 | 0.23 | 0.72 |
| Taxidea taxus | semifossorial | semifossorial | 0 | 0 | 0 | 0.79 | 0.21 |
| Bassaricyon alleni | arboreal | arboreal | 0.99 | 0 | 0.01 | 0 | 0 |
| Bassaricyon gabbii | arboreal | arboreal | 1 | 0 | 0 | 0 | 0 |
| Bassaricyon medius | arboreal | arboreal | 1 | 0 | 0 | 0 | 0 |
| Bassaricyon neblina | arboreal | arboreal | 0.87 | 0 | 0.13 | 0 | 0 |
| Nasua narica | scansorial | scansorial | 0.05 | 0 | 0.93 | 0 | 0.02 |
| Nasuella olivacea | scansorial | scansorial | 0.46 | 0 | 0.47 | 0 | 0.07 |
| Nasua nasua | scansorial | scansorial | 0 | 0 | 0.81 | 0.01 | 0.18 |
| Bassariscus astutus | scansorial | scansorial | 0.46 | 0 | 0.52 | 0 | 0.02 |
| Procyon cancrivorus | scansorial | scansorial | 0.01 | 0 | 0.87 | 0 | 0.12 |
| Procyon lotor | scansorial | scansorial | 0.03 | 0 | 0.88 | 0 | 0.09 |
| Potos flavus | arboreal | arboreal | 0.96 | 0 | 0.04 | 0 | 0 |
| Conepatus mesoleucus | terrestrial | terrestrial | 0 | 0 | 0.12 | 0.07 | 0.81 |
| Mephitis mephitis | terrestrial | terrestrial | 0 | 0 | 0.05 | 0.19 | 0.76 |
| Spilogale gracilis | terrestrial | terrestrial | 0 | 0 | 0.39 | 0.02 | 0.59 |
| Mydaus javanensis | terrestrial | terrestrial | 0 | 0 | 0.51 | 0.26 | 0.22 |
| Atelocynus microtis | terrestrial | terrestrial | 0 | 0 | 0.16 | 0.09 | 0.75 |
| Cerdocyon thous | terrestrial | terrestrial | 0 | 0 | 0.2 | 0 | 0.79 |
| Lycalopex griseus | terrestrial | terrestrial | 0 | 0 | 0.04 | 0 | 0.96 |
| Chrysocyon brachyurus | terrestrial | terrestrial | 0 | 0 | 0.56 | 0 | 0.44 |
| Speothos venaticus | terrestrial | semifossorial | 0 | 0 | 0 | 0.85 | 0.15 |
| Canis adustus | terrestrial | terrestrial | 0 | 0 | 0 | 0 | 0.99 |
| Canis mesomelas | terrestrial | terrestrial | 0 | 0 | 0.03 | 0 | 0.97 |
| Canis aureus | terrestrial | terrestrial | 0 | 0 | 0.01 | 0 | 0.99 |
| Canis latrans | terrestrial | terrestrial | 0 | 0 | 0.07 | 0 | 0.93 |
| Canis lupus | terrestrial | terrestrial | 0 | 0 | 0.03 | 0.04 | 0.93 |
| Cuon alpinus | terrestrial | terrestrial | 0 | 0 | 0.33 | 0.02 | 0.65 |
| Lycaon pictus | terrestrial | terrestrial | 0 | 0 | 0.02 | 0 | 0.98 |
| Nyctereutes procyonoides | terrestrial | terrestrial | 0 | 0 | 0.56 | 0.01 | 0.43 |
| Urocyon cinereoargenteus | scansorial | terrestrial | 0 | 0 | 0.07 | 0.01 | 0.92 |
| Vulpes zerda | terrestrial | terrestrial | 0 | 0 | 0.01 | 0 | 0.99 |
| Vulpes rueppellii | terrestrial | terrestrial | 0 | 0 | 0 | 0 | 1 |
| Vulpes vulpes | terrestrial | terrestrial | 0 | 0 | 0.13 | 0 | 0.87 |
| Vulpes lagopus | terrestrial | terrestrial | 0 | 0 | 0 | 0 | 1 |
| Otocyon megalotis | terrestrial | terrestrial | 0 | 0 | 0.01 | 0 | 0.99 |
