## Supplementary FIle B for "Topographically distinct adaptive landscapes for teeth, skeletons, and size explain the adaptive radiation of Carnivora (Mammalia)"

### Supplementary File B: Carnivoran post-cranial ratio analyses

In this document we'll assess the fit of the canonical three macroevolutionary models to each of the carnivoran post-cranial ratios (functional indices). Support for an early burst model over all other models would indicate early rapid evolution of this axis of morphological variation. Support for a single optimum OU model (single stationary peak) is consistent with constraint (high alpha) or late accumulation of variation (low alpha). Support for Brownian motion over the other models should never be high but would be consistent with evolution at a relatively constant rate, adaptive evolution chasing a fluctuating optimum, among other options.

First, we will load required packages and some supplementary functions

```
source("functions_postcranial.R")
library(geiger)
```

```
## Loading required package: ape
```

Now we will read in the required data: 1) a time-scaled phylogeny of carnivorans from Slater and Friscia 2019; 2) a csv file of species mean ratio values, and; 3) a csv file of associated standard errors. These files are in the postcranial_ratios directory. After doing this, we will prune un-needed taxa from the tree

```
phy<-ladderize(read.tree("postcranial_ratios/mcc.tre"))
mean.rat <- read.csv("postcranial_ratios/mean.ratios.csv", row.names=1)
sems <- read.csv("postcranial_ratios/sems.csv", row.names=1)
phy<-treedata(phy, mean.rat,warnings = F)$phy
```

We will now use a set of loops to fit the three models to each column of the matrix. By doing this, we can store the output in a list to later extract AICc scores and parameter estimates.

```
bm.fit<- list()
for(i in 1:ncol(mean.rat)) {
  bm.fit[[i]] <- fitContinuous(phy=phy, dat=setNames(mean.rat[,i],
                 rownames(mean.rat)), SE=setNames(sems[,i], rownames(sems)), model="BM")
}

eb.fit<- list()
for(i in 1:ncol(mean.rat)) {
  eb.fit[[i]] <- fitContinuous(phy=phy, dat=setNames(mean.rat[,i],
                rownames(mean.rat)), SE=setNames(sems[,i], rownames(sems)), model="EB")
}
```

```
## Warning in fitContinuous(phy = phy, dat = setNames(mean.rat[, i], rownames(mean.rat)), :
## Parameter estimates appear at bounds:
##  a

## Warning in fitContinuous(phy = phy, dat = setNames(mean.rat[, i], rownames(mean.rat)), :
## Parameter estimates appear at bounds:
##  a

## Warning in fitContinuous(phy = phy, dat = setNames(mean.rat[, i], rownames(mean.rat)), :
## Parameter estimates appear at bounds:
##  a
```

```
## Warning in fitContinuous(phy = phy, dat = setNames(mean.rat[, i], rownames(mean.rat)), :
## Parameter estimates appear at bounds:
##  a

## Warning in fitContinuous(phy = phy, dat = setNames(mean.rat[, i], rownames(mean.rat)), :
## Parameter estimates appear at bounds:
##  a

## Warning in fitContinuous(phy = phy, dat = setNames(mean.rat[, i], rownames(mean.rat)), :
## Parameter estimates appear at bounds:
##  a

## Warning in fitContinuous(phy = phy, dat = setNames(mean.rat[, i], rownames(mean.rat)), :
## Parameter estimates appear at bounds:
##  a

## Warning in fitContinuous(phy = phy, dat = setNames(mean.rat[, i], rownames(mean.rat)), :
## Parameter estimates appear at bounds:
##  a

## Warning in fitContinuous(phy = phy, dat = setNames(mean.rat[, i], rownames(mean.rat)), :
## Parameter estimates appear at bounds:
##  a

## Warning in fitContinuous(phy = phy, dat = setNames(mean.rat[, i], rownames(mean.rat)), :
## Parameter estimates appear at bounds:
##  a

## Warning in fitContinuous(phy = phy, dat = setNames(mean.rat[, i], rownames(mean.rat)), :
## Parameter estimates appear at bounds:
##  a

## Warning in fitContinuous(phy = phy, dat = setNames(mean.rat[, i], rownames(mean.rat)), :
## Parameter estimates appear at bounds:
##  a

## Warning in fitContinuous(phy = phy, dat = setNames(mean.rat[, i], rownames(mean.rat)), :
## Parameter estimates appear at bounds:
##  a

## Warning in fitContinuous(phy = phy, dat = setNames(mean.rat[, i], rownames(mean.rat)), :
## Parameter estimates appear at bounds:
##  a

## Warning in fitContinuous(phy = phy, dat = setNames(mean.rat[, i], rownames(mean.rat)), :
## Parameter estimates appear at bounds:
##  a
```

```r
ssp.fit<- list()
for(i in 1:ncol(mean.rat)) {
  ssp.fit[[i]] <- fitContinuous(phy=phy, dat=setNames(mean.rat[,i],
                  rownames(mean.rat)), SE=setNames(sems[,i], rownames(sems)), model="OU")
}
```

Having done this, we still need to perform model selection to chose the best-fitting model for each trait. We will first create a function to do this, and a matrix to hold the output (AIC weights for each model):

```
compare.models <- function(x,y,z) {
  geiger:::aicw(c(x$opt$aicc, y$opt$aicc, z$opt$aicc))[,3]
}
res<-matrix(nrow=ncol(mean.rat), ncol=3)
```

Now we'll loop through and, for each trait, compute AIC weights for each model

```
for(i in 1:length(bm.fit)) {
  res[i,]<-compare.models(bm.fit[[i]], ssp.fit[[i]], eb.fit[[i]])
}
colnames(res)<- c("BM", "SSP", "EB"); rownames(res) <- colnames(mean.rat)
```

We can look at these results in table form

```
##                               BM  SSP   EB
## scapula.index               0.59 0.21 0.21
## glenoid.shape               0.01 0.99 0.00
## brachial.index              0.12 0.83 0.04
## humeral.epicondylar.breadth 0.52 0.29 0.18
## capitulum.shape             0.38 0.49 0.13
## fossoriality.index          0.16 0.78 0.06
## crural.index                0.37 0.50 0.13
## femoral.shaft.shape         0.00 1.00 0.00
## femoral.epicondylar.width   0.52 0.30 0.18
## patella.grove.index         0.44 0.40 0.16
## femoral.epicondylar.index   0.23 0.69 0.08
## gluteal.index               0.44 0.40 0.16
## intermembranal.index        0.39 0.47 0.14
## ischial.breadth             0.12 0.83 0.04
## iliac.breadth               0.58 0.21 0.20
## pubic.symphysis.length      0.57 0.23 0.20
```

or visualize them as stacked bar-plots

```
par(mar=c(5,5,2,5), mfrow=c(1,1), xpd=T)

barplot(t(res), col=c("yellow","skyblue", "black"), space = 0, las=1,
        cex.names = 1, ylab="Akaike Weight", names.arg = seq(1,16), xlab = "Functional Trait")

legend(x=16.25,y=0.5, legend = c("BM", "SSP", "EB"),
      pch = 22,
       pt.bg =c("yellow", "lightblue", "black"),
       pt.cex=2, cex = 1)
```

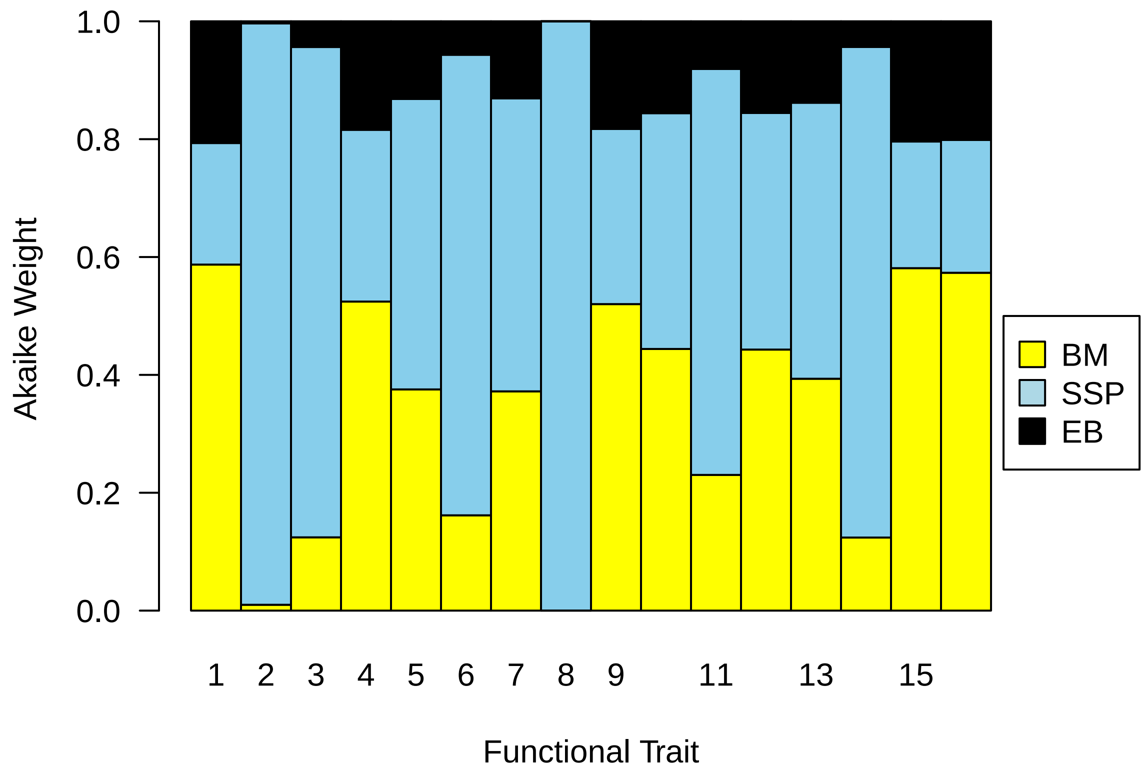


Both forms of output are saved in the ratio analysis folder.

Finally, we might like to look at the parameter estimates for each best-fit model to all nuanced interpretation of macroevolutionary dynamics. We can do so by querying the res list

```
best.model <- apply(res, 1, which.max)
best.model.params <- matrix(data=NA, nrow=16, ncol=5)
colnames(best.model.params) <- c("Model", "Weight", "Root", "Rate", expression(alpha))
rownames(best.model.params) <- colnames(mean.rat)
for(i in 1:length(best.model)) {
  if(best.model[i]==1) best.model.params[i,]<- c("BM", round(res[i,1],2), round(bm.fit[[i]]$opt$z0,2), b
  if(best.model[i]==2) best.model.params[i,]<- c("SSP", round(res[i,2],2),round(ssp.fit[[i]]$opt$z0,2),s
}

best.model.params
```

```
##                             Model Weight Root   Rate                    alpha
## scapula.index               "BM"  "0.59" "0.85" "0.000286015127985488" "-"
## glenoid.shape               "SSP" "0.99" "0.69" "0.000210788885057432" "0.05"
## brachial.index              "SSP" "0.83" "0.84" "0.000632718344441387" "0.03"
## humeral.epicondylar.breadth "BM"  "0.52" "0.25" "9.59420697740497e-05" "-"
## capitulum.shape             "SSP" "0.49" "1.46" "0.000493540950044172" "0.03"
## fossoriality.index          "SSP" "0.78" "0.14" "7.11651568480824e-05" "0.03"
## crural.index                "SSP" "0.5"  "0.93" "0.000518587202687116" "0.02"
## femoral.shaft.shape         "SSP" "1"    "1.12" "0.00101647889026685"  "0.11"
## femoral.epicondylar.width   "BM"  "0.52" "0.19" "2.72898358839809e-05" "-"
## patella.grove.index         "BM"  "0.44" "0.43" "6.57548891403957e-05" "-"
## femoral.epicondylar.index   "SSP" "0.69" "0.01" "1.46798699158029e-06" "0.03"
## gluteal.index               "BM"  "0.44" "0.18" "2.8843462167798e-05"  "-"
## intermembranal.index        "SSP" "0.47" "0.84" "0.00019923058153718"  "0.02"
```

```
## ischial.breadth             "SSP" "0.83" "0.58" "0.000461712247461033" "0.03"
## iliac.breadth               "BM"  "0.58" "0.64" "0.000602946526330765" "-"
## pubic.symphysis.length      "BM"  "0.57" "0.27" "0.000145508407003235" "-"
```

In no case is an EB model recovered as best-fitting. Most cases in which an SSP model are preferred yield relatively low alpha values, corresponding to phylogenetic half-lives of 13-23 myr (crown carnivora is ~ 36 myr old). The exception is femoral shape shape, which yields an alpha of 0.11 and half-life of 6.3 myr.
