## Supplementary File D for "Topographically distinct adaptive landscapes for teeth, skeletons, and size explain the adaptive radiation of Carnivora (Mammalia)"

### Supplementary File D: Center of Gravity Simulations

In this document we will do some simple simulations to confirm that the COG for a univariate trait evolving at a constant rate on a molecular phylogeny is ~ 0.5, and then that the COG for a trait evolving under a strict Early Burst (declining rates) is lower than 0.5

```
library(phytools)
```

```
## Loading required package: ape
```

```
## Loading required package: maps
```

```
library(geiger)
library(TreeSim)
```

First, define a matrix to hold our results

```
results <- matrix(nrow=1000, ncol=3)
```

Now we'll simulate Yule trees and BM evolution, computing dtt, CoG and the gamma statistic, each time

```
for(i in 1:nrow(results)) {
  phy <- pbtree(n=100)
  d <- fastBM(phy)
  disp <- dtt(phy, d, plot = F)
  CoG<-sum(disp$dtt * disp$times) / sum(disp$dtt)
  gs <- gammaStat(phy)
  root.age <- max(diag(vcv(phy)))
  results[i,] <- c(root.age, gs, CoG)
}
```

First, let's look at the distribution of COGs

```
hist(results[,3], col="dodgerblue", xlab="Center of Gravity", main="center of gravity on Yule trees")
```

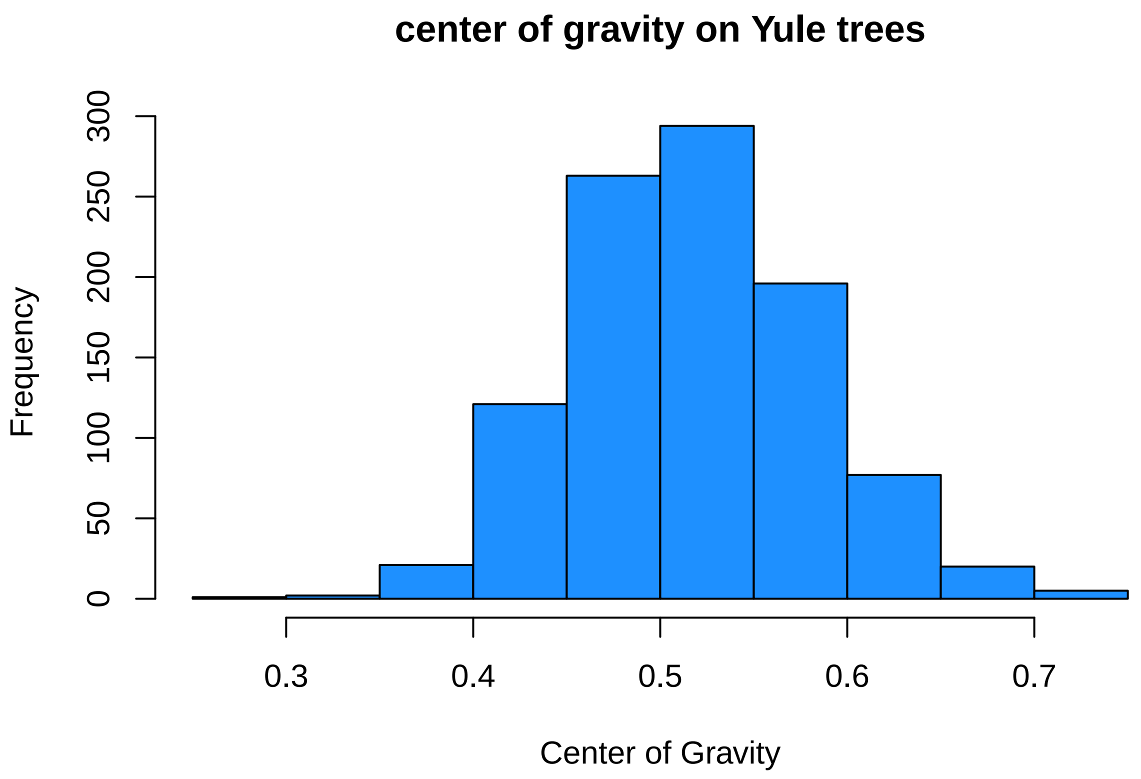


```
mean(results[,3])
```

```
## [1] 0.5174119
```

```
quantile(results[,3], c(0.025,0.975))
```

```
##      2.5%     97.5%
## 0.4009884 0.6491396
```

So the CoG is essentially 0.5. Is there a relationship betweeen the distribution of branching events and the CoG?

```
mymodel <- lm(results[,3]~results[,2])
summary(mymodel)
```

```
##
## Call:
## lm(formula = results[, 3] ~ results[, 2])
##
## Residuals:
##       Min        1Q    Median        3Q       Max
## -0.243338 -0.041140 -0.001185  0.040562  0.216973
##
## Coefficients:
##              Estimate Std. Error t value Pr(>|t|)
## (Intercept)  0.517323   0.001997 259.020  < 2e-16 ***
## results[, 2] 0.008479   0.001937   4.377 1.33e-05 ***
## ---
## Signif. codes:  0 '***' 0.001 '**' 0.01 '*' 0.05 '.' 0.1 ' ' 1
##
## Residual standard error: 0.06315 on 998 degrees of freedom
## Multiple R-squared:  0.01883,    Adjusted R-squared:  0.01785
```

```
## F-statistic: 19.16 on 1 and 998 DF,  p-value: 1.331e-05
```

```r
plot(results[,2:3], xlab="gamma", ylab="CoG", pch=21, bg="dodgerblue")
abline(summary(mymodel))
```

```
## Warning in abline(summary(mymodel)): only using the first two of 8 regression
## coefficients
```

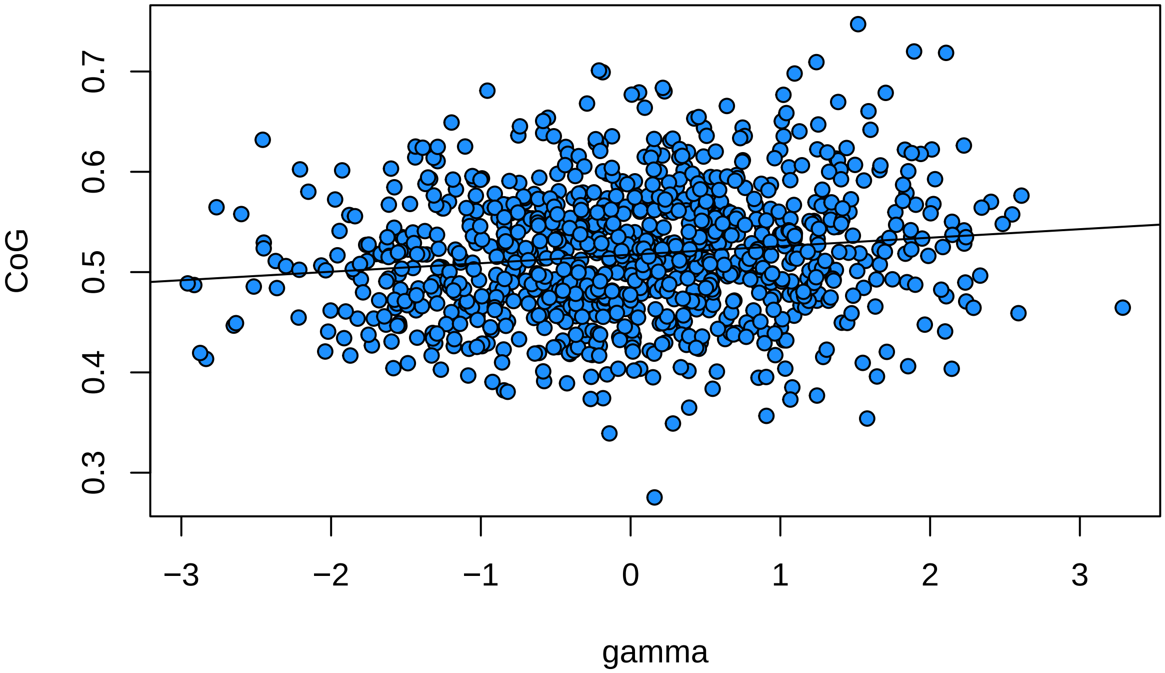


So yes, a little bit - more tippy trees (more positive gamma) have slightly higher CoGs - but only 1.5% of the variance in the COG is associated with the gamma statistic. We can double check by simulating under a birth death process with high extinction, generating really tippy trees.

```r
resultsBD <- matrix(nrow=1000, ncol=3)

for(i in 1:nrow(resultsBD)) {
  phy <- sim.bd.taxa(n=100, 1, lambda=1, mu=0.95, complete=F)[[1]]
  d <- fastBM(phy)
  disp <- dtt(phy, d, plot = F)
  CoG<-sum(disp$dtt * disp$times) / sum(disp$dtt)
  gs <- gammaStat(phy)
  root.age <- max(diag(vcv(phy)))
  resultsBD[i,] <- c(root.age, gs, CoG)
}
```

```r
hist(resultsBD[,3], col="dodgerblue", xlab="Center of Gravity", main="center of gravity on Yule trees")
```

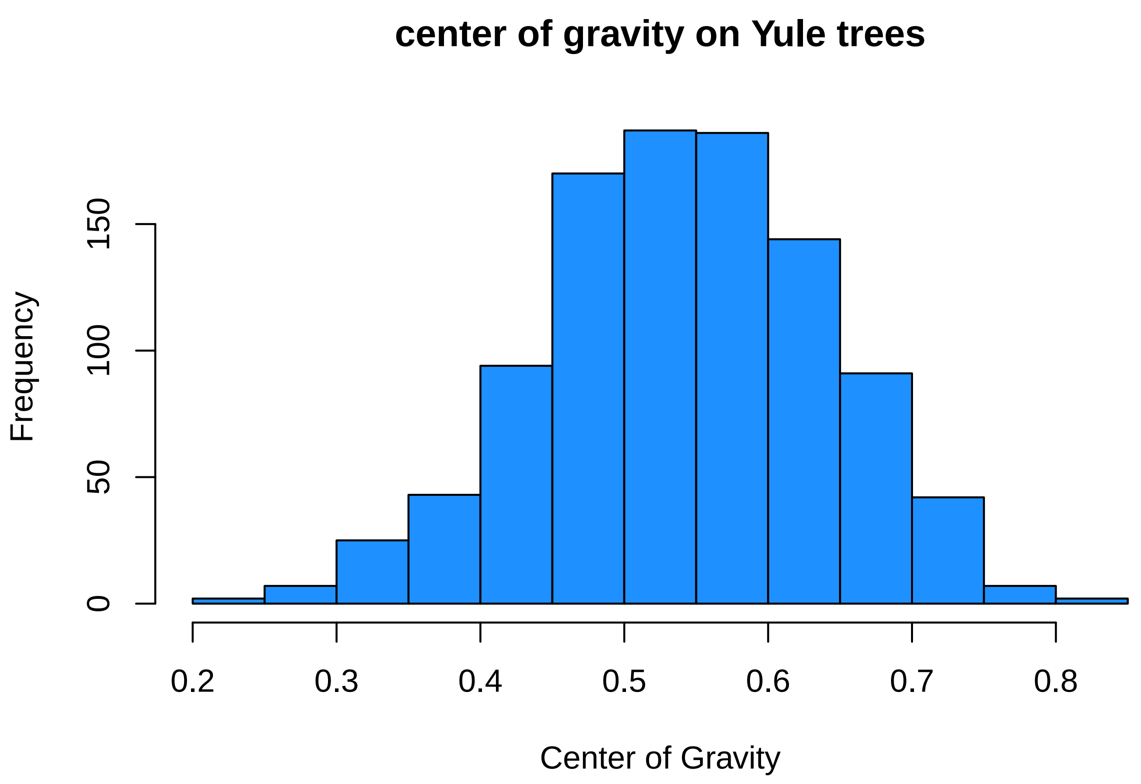


```
mean(resultsBD[,3])
```

```
## [1] 0.541115
```

```
quantile(resultsBD[,3], c(0.025,0.975))
```

```
##      2.5%     97.5%
## 0.3283491 0.7290816
```

```
mymodel <- lm(resultsBD[,3]~resultsBD[,2])
summary(mymodel)
```

```
##
## Call:
## lm(formula = resultsBD[, 3] ~ resultsBD[, 2])
##
## Residuals:
##       Min        1Q    Median        3Q       Max
## -0.315813 -0.068522  0.002135  0.069411  0.288051
##
## Coefficients:
##                  Estimate Std. Error t value Pr(>|t|)
## (Intercept)     0.606494   0.015036  40.337  < 2e-16 ***
## resultsBD[, 2] -0.009358   0.002106  -4.444  9.8e-06 ***
## ---
## Signif. codes:  0 '***' 0.001 '**' 0.01 '*' 0.05 '.' 0.1 ' ' 1
##
## Residual standard error: 0.09836 on 998 degrees of freedom
## Multiple R-squared:  0.01941,    Adjusted R-squared:  0.01842
## F-statistic: 19.75 on 1 and 998 DF,  p-value: 9.804e-06
```

```
plot(resultsBD[,2:3], xlab="gamma", ylab="CoG", pch=21, bg="dodgerblue")
abline(summary(mymodel))
```

```
## Warning in abline(summary(mymodel)): only using the first two of 8 regression
## coefficients
```

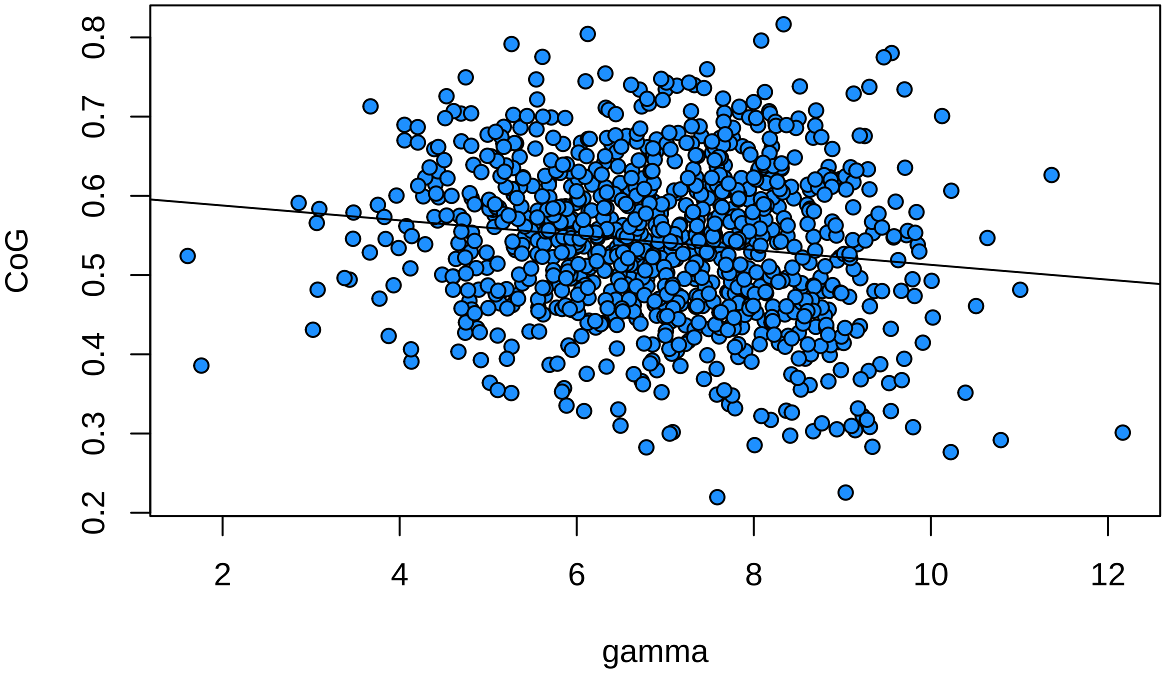


Interestingly, the mean CoG here moves upwards (as do the 95% quantiles) but the relationship between gamma and CoG is actually negative over this range of topologies and branch lengths. It seems likely that for the tippiest trees, the COG will be low because extinction will have pruned out enough taxa to create a set of young crown clades, whithin which members share more similar traits than they do among clades. This emphasizes the importance of assessing significance on the topology in question.

```
  bd_sig_res <- matrix(data=NA, nrow=1000, ncol=2)
  for(i in 1:1000) {
    phy <- sim.bd.taxa(n=100, 1, lambda=1, mu=0.95, complete=F)[[1]]
    d <- fastBM(phy)
    disp <- dtt(phy, d, nsim = 999, plot = F)
    CoG<-sum(disp$dtt * disp$times) / sum(disp$dtt)
    sim.cog<-apply(disp$sim, 2, function(x) sum(x*disp$times) / sum(x))
    p<-length(which(sim.cog<CoG)) / (length(sim.cog))
    bd_sig_res[i, ]<-c(CoG, p)
  }
```

let's look at the type 1 error rate

```
hist(bd_sig_res[,2], col="dodgerblue", breaks=100, xlab="p value", main=paste("false positive rate = ",
abline(v=0.05, col="red", lty=2, lwd=5)
```

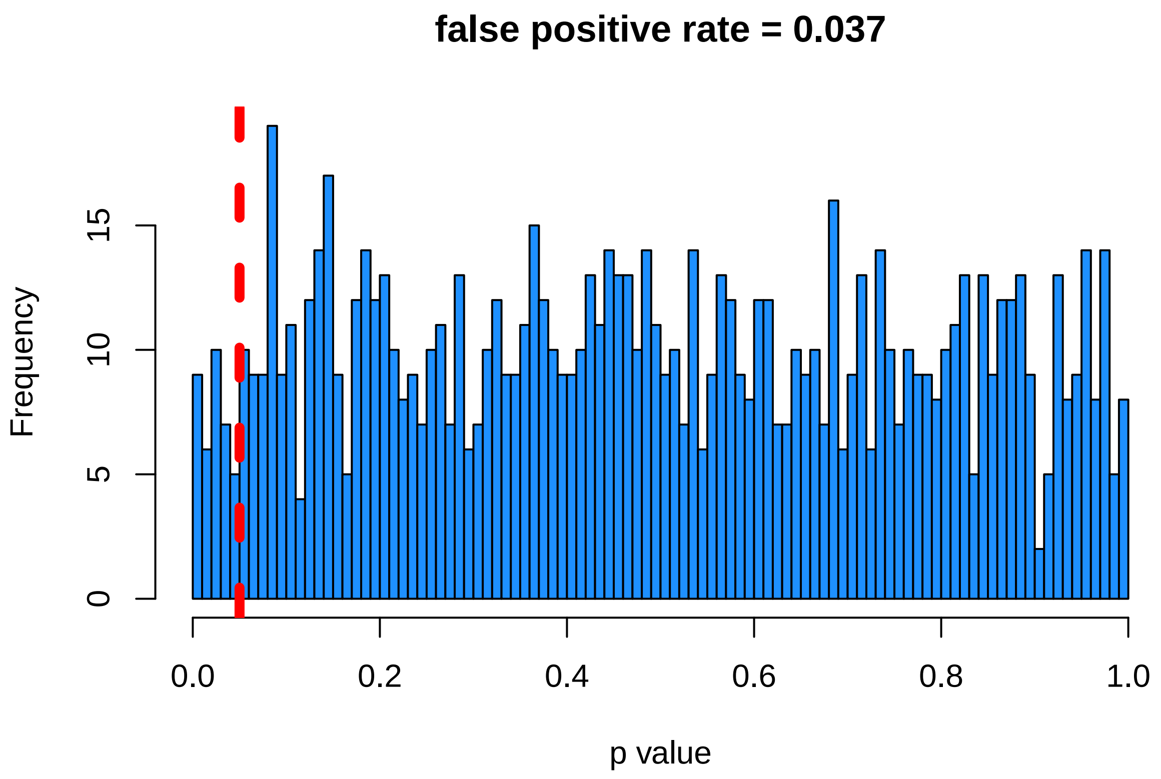


looks ok - it seems as though we have an acceptable rate of falsely rejected the true constant rates model here.

##### Early Bursts

We can now ask how we do at identifying early bursts - evolutionary rates that slow through time. Let's start out with the Yule process like before. We'll set it up so that rates decline 7 half lifes over clade history, which is enough to leave a strong signal (see Slater and Pennell Syst Biol 2014)

```
results <- matrix(nrow=1000, ncol=3)

for(i in 1:nrow(results)) {
  phy <- pbtree(n=100)
  root.age <- max(diag(vcv(phy)))
  r <- log(2)/(root.age/7)
  eb.phy <- rescale(phy, model = "EB", -r)
  d <- fastBM(eb.phy)
  disp <- dtt(phy, d, plot = F)
  CoG<-sum(disp$dtt * disp$times) / sum(disp$dtt)
  gs <- gammaStat(phy)
  root.age <- max(diag(vcv(phy)))
  results[i,] <- c(root.age, gs, CoG)
}
hist(results[,3], col="dodgerblue", xlab="Center of Gravity", main="center of gravity on Yule trees unde
```

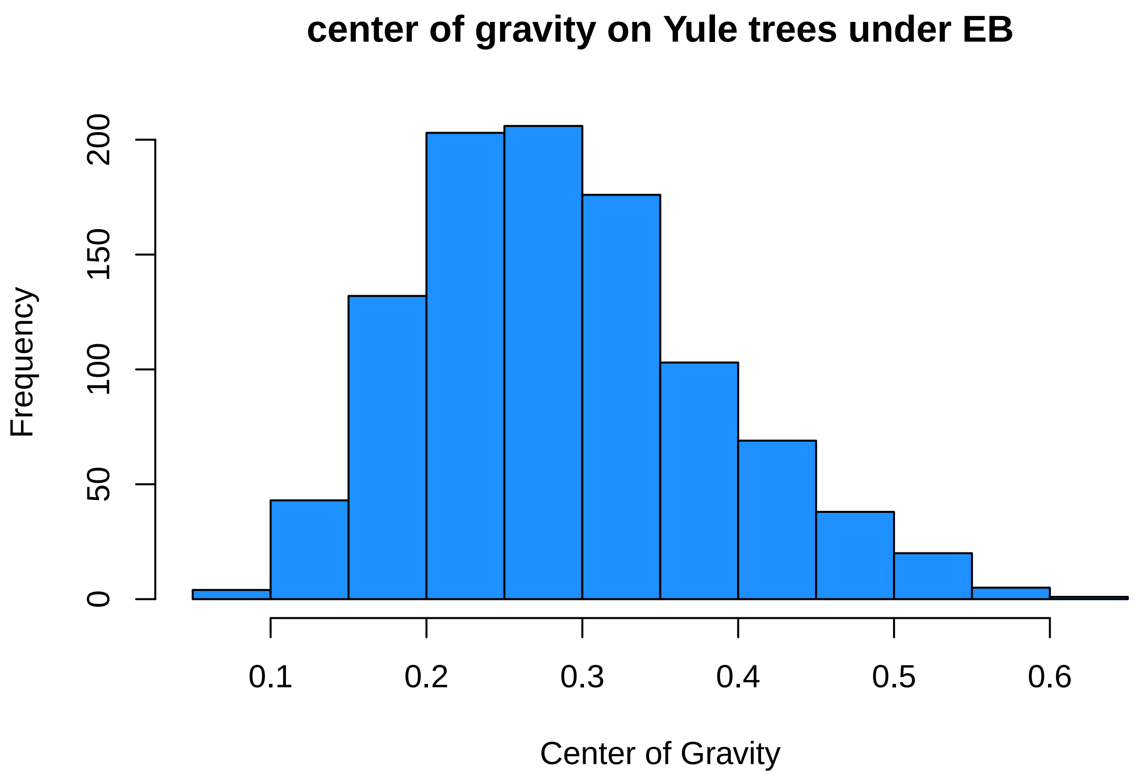


```
mean(results[,3])
```

```
## [1] 0.2876862
```

```
quantile(results[,3], c(0.025,0.975))
```

```
##      2.5%     97.5%
## 0.1348489 0.5004140
```

The CoG is negatively shifted here and the 95% quantiles do not include 0.5. This is the kind of situation we would hope for. But, we should be more concerned about cases in which the true tree is a bd tree as exinction happens, so lets look at that:

```
results <- matrix(nrow=1000, ncol=2)

for(i in 1:nrow(results)) {
    phy <- sim.bd.taxa(n=100, 1, lambda=1, mu=0.95, complete=F)[[1]]
    root.age <- max(diag(vcv(phy)))
    r <- log(2)/(root.age/7)
    eb.phy <- rescale(phy, model = "EB", -r)
    d <- fastBM(eb.phy)
    disp <- dtt(phy, d, nsim = 999, plot = F)
    CoG<-sum(disp$dtt * disp$times) / sum(disp$dtt)
    sim.cog<-apply(disp$sim, 2, function(x) sum(x*disp$times) / sum(x))
    p<-length(which(sim.cog<CoG)) / (length(sim.cog))
    bd_sig_res[i, ]<-c(CoG, p)
}
```

```
hist(bd_sig_res[,1], col="dodgerblue", xlab="Center of Gravity", main="center of gravity on BD trees und
```

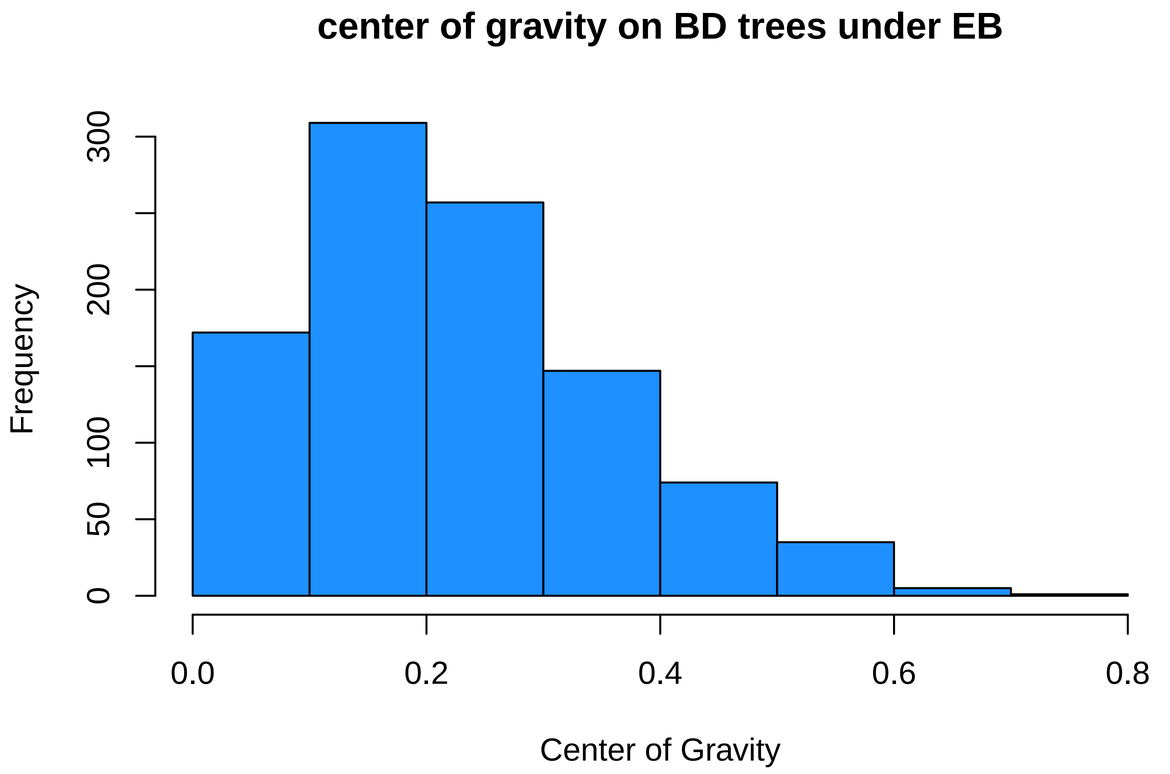


```
mean(bd_sig_res[,1])
```

```
## [1] 0.2279038
```

```
quantile(bd_sig_res[,1], c(0.025,0.975))
```

```
##      2.5%     97.5%
## 0.0458215 0.5433649
```

The mean is a little lower (0.23 vs 0.28), but the quantiles are larger (0.04-0.55 vs 0.13-0.49). We can look at how this affects power - our ability to reject the null:

```
hist(bd_sig_res[,2], col="dodgerblue", breaks=100, xlab="p value", main=paste("power = ", round(length(
abline(v=0.05, col="red", lty=2, lwd=5)
```

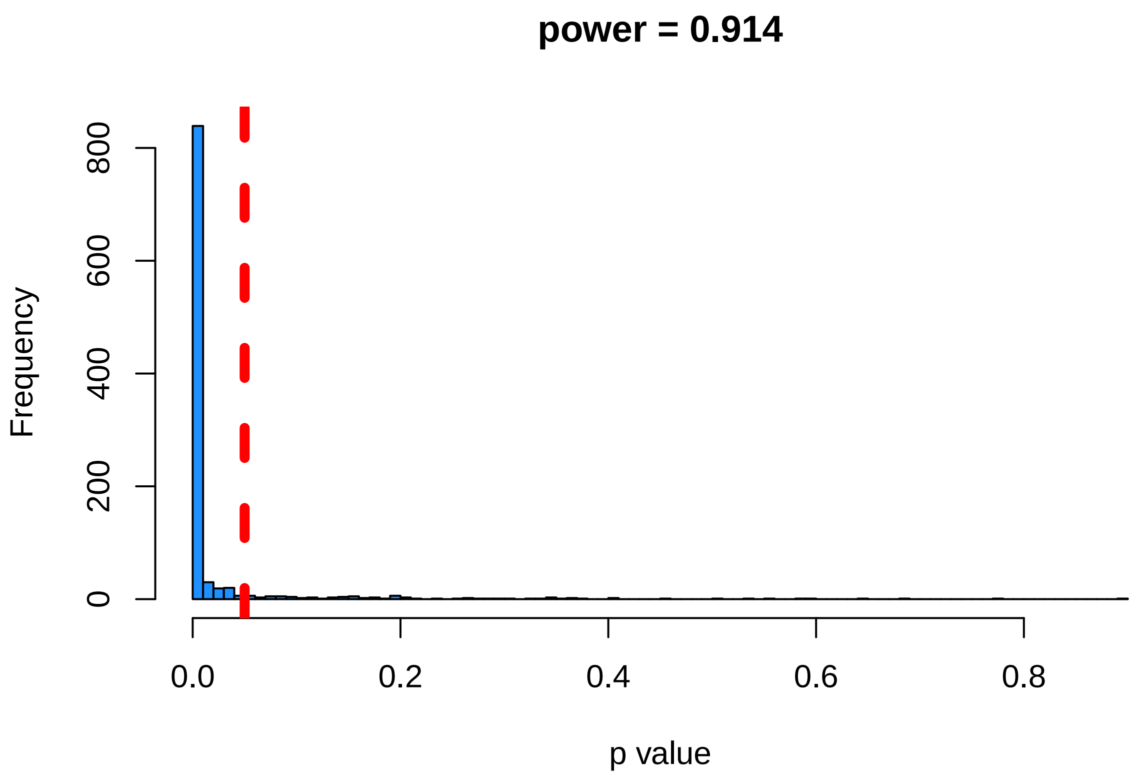


Looks like we have lots of power to detect an early burst when it happens, and our type I (false positive) rate is similarly low. Overall, the DTT Center of Gravity seens like an appropriate metric to test for "early burst-like" dynamics in comparative data.
